# Tissue-specific RNA-seq defines genes governing male tail tip morphogenesis in *C. elegans*

**DOI:** 10.1101/2024.01.12.575210

**Authors:** Karin Kiontke, R. Antonio Herrera, D. Adam Mason, Alyssa Woronik, Stephanie Vernooy, Yash Patel, David H. A. Fitch

## Abstract

*Caenorhabditis elegans* males undergo sex-specific tail tip morphogenesis (TTM) under the control of the transcription factor DMD-3. To find genes regulated by DMD-3, We performed RNA-seq of laser-dissected tail tips. We identified 564 genes differentially expressed (DE) in wild-type males vs. *dmd-3(-)* males and hermaphrodites. The transcription profile of *dmd-3(-)* tail tips is similar to that in hermaphrodites. For validation, we analyzed transcriptional reporters for 49 genes and found male-specific or male-biased expression for 26 genes. Only 11 DE genes overlapped with genes found in a previous RNAi screen for defective TTM. GO enrichment analysis of DE genes finds upregulation of genes within the UPR (unfolded protein response) pathway and downregulation of genes involved in cuticle maintenance. Of the DE genes, 40 are transcription factors, indicating that the gene network downstream of DMD-3 is complex and potentially modular. We propose modules of genes that act together in TTM and are coregulated by DMD-3, among them the chondroitin synthesis pathway and the hypertonic stress response.

## Introduction

Cellular morphogenesis, a process essential for animal development, involves precise temporal, spatial and often sex-specific coordination of various cytological events, such as cell fusion, change in cell shape and migration. It has been studied mostly in embryonic processes like tubulogenesis, gastrulation, neural crest migration and the epithelial-to-mesenchymal transition (EMT) (Debnath et al., 2022; Gheisari et al., 2020; Goldstein and Nance, 2020; Hashimoto and Munro, 2019; Iruela-Arispe and Beitel, 2013; Shaye and Soto, 2021; Szabó and Mayor, 2018). However, it is also important for postembryonic structures, such as the excretory system of *C. elegans*, a model for tubulogenesis (Shaye and Soto, 2021), and sexual dimorphisms manifested during the juvenile-to-adult transition in animals (Kopp, 2011; Mason et al., 2008; Ohde et al., 2018). In *C. elegans,* sexually dimorphic morphogenesis forms the vulva and uterus of hermaphrodites (Gupta et al., 2012) and the male tail (Mason et al., 2008; Nelson et al., 2011; Nguyen et al., 1999). In nearly all cases, a major deficiency in our understanding is how the cell biological events of morphogenesis are regulated transcriptionally in the gene regulatory network (GRN).

To investigate the transcriptional regulation of sexually dimorphic morphogenesis, we focus here on the male tail of *C. elegans*. In larvae, the male tail is morphologically similar to that of hermaphrodites and has a pointed tip (Fig. 1). In adult males, however, the tail tip is short and round. The hermaphrodite tail tip retains the long, pointed shape of the juvenile. The morphogenetic process leading to the short and round male tail tip happens at 4th larval stage (L4) during the juvenile-to-adult transition and is called male tail tip morphogenesis (TTM).

**Figure 1.**
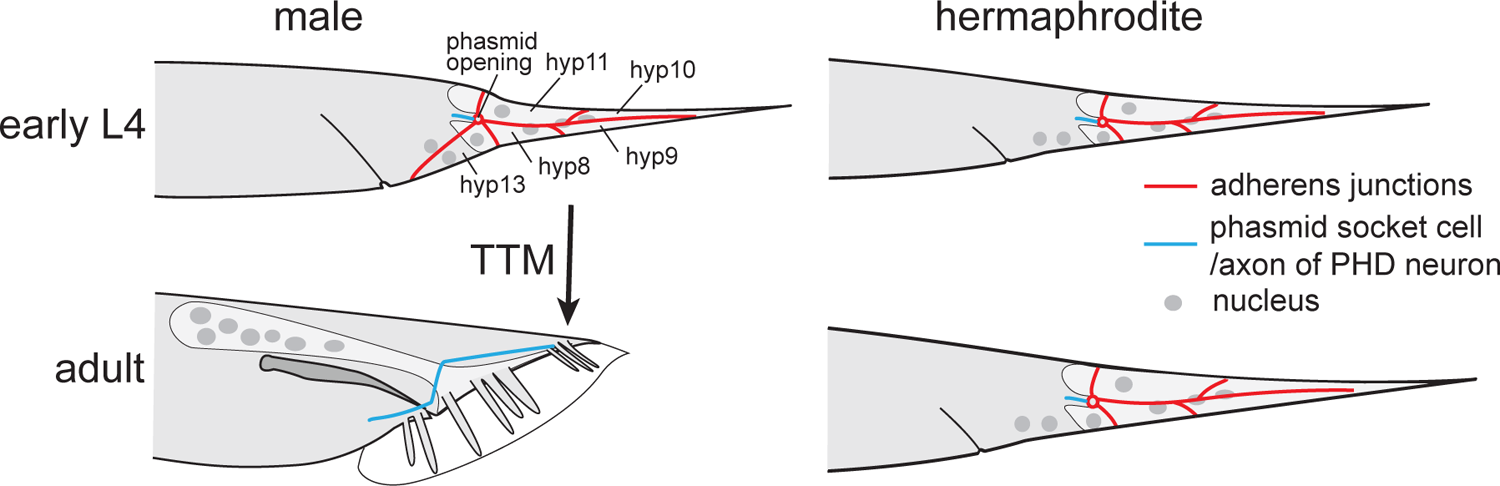
Tail tips in *C. elegans*. The larvae of both sexes have a long, pointed tail consisting of 4 epithelial cells, hyp8-11. Hermaphrodites retain this shape as adults. In males, the morphogenetic process TTM creates a short and round tail. Adherens junctions between the tail tip cells disassemble. One cell of each phasmid socket transdifferentiates into the male-specific PHD neuron.

During this time, other male-specific structures form in the tail: a pair of spicules and 9 pairs of sensory processes called rays that are embedded in a cuticular fan or bursa. TTM in *C. elegans* was first described by Sulston et al. (1980), who provided a detailed map of the tail nuclei at four time points during TTM. Later, Nguyen et al. (1999) described the shape change of the tail tip cells at the beginning of TTM at the ultrastructural level. These studies showed that the tail tip of males and hermaphrodites consists of 4 cells, hyp 8-11, that originate during embryogenesis. In males only, an additional binucleate cell, hyp13, is present ventrally just anterior of the tail tip. In males, at the beginning of the L4 stage, the adherens junctions (AJ) between hyp8-11 disassemble and the cells begin to fuse, although they initially retain much of the plasma membranes between them. Subsequently, the tail tip tissue rounds up and shortens.

Neurons in the tail also undergo sexually dimorphic morphogenesis: In L3 males and hermaphrodites, two PHC neurons send a dendritic process into the tip of the tail (hyp10) and an axon into the preanal ganglion (White et al., 1986). In males only, during L4, the PHC dendritic process shortens during TTM, and the axon grows out anteriorly and differentiates into a male-specific hub neuron that is required for male mating behavior (Serrano-Saiz et al., 2017). A second pair of male-specific neurons, PHDL and PHDR, develop de-novo through transdifferentiation and reorganization of the phasmid socket cells PHso1L and PHso1R (Molina-García et al., 2020).

Previous research (Mason et al., 2008) demonstrated that TTM is controlled by the DM-domain transcription factor DMD-3, a homolog of *Drosophila* Doublesex and mammalian Dmrt1. DMD-3 is male-specifically expressed and both necessary and sufficient for TTM. Loss-of-function mutations in the *dmd-3* gene result in partial failure of TTM, and misexpression of DMD-3 in hermaphrodites leads to ectopic TTM. TTM, and indeed all male tail morphogenesis, fails completely in double loss-of-function mutants of *dmd-3* and its paralog *mab-3*. However, *mab-3* loss-of-function mutants have low-penetrance TTM phenotypes (Shen and Hodgkin, 1988), suggesting *mab-3* contributes to robustness of TTM but that *dmd-3* has the major role as a “master regulator” of the process. *dmd-3* is also necessary and sufficient for the differentiation of the PHC neurons into male hub neurons (Serrano-Saiz et al., 2017).

A genome-wide RNAi screen for TTM defects found 210 genes that contribute to TTM (Nelson et al., 2011). Many of these genes play a regulatory role and belong to pathways that determine, for example, when (via the heterochronic pathway), where (via HOX transcription factors) and in which sex (via the sex determination pathway) TTM takes place. Other genes are involved in the processes that execute TTM, e.g. genes involved in fusion, vesicular transport or rearrangement of the cytoskeleton. The current hypothesis for the gene network for TTM is that it has a bow-tie architecture with DMD-3 and MAB-3 at the core (Nelson et al., 2011).

Regulatory pathways determine the expression of DMD-3, which then coordinates the cell biological processes during morphogenesis. How DMD-3 is linked to these cell biological processes is insufficiently known. In fact, how transcriptional regulators communicate with the cell machinery to drive morphogenesis is largely unknown for most systems (Bernadskaya and Christiaen, 2016; Debnath et al., 2022; Gildor et al., 2021; Kenny-Ganzert and Sherwood, 2024; Shaye and Soto, 2021; Yamakawa et al., 2023).

To address this question, we sought to identify genes that are specifically expressed in the tail tip during TTM. Our strategy was to compare the transcriptomes of tail tips that undergo TTM (wild-type males and *dmd-3(gf)* hermaphrodites ectopically expressing DMD-3) to the transcriptomes of tail tips that do not (wild-type hermaphrodites and *dmd-3(lf)* males). Because the tail tip is at the end of the animal, it can easily be isolated using laser capture microdissection. A correctly placed cut can isolate the 4 tail tip cells at the beginning of TTM, which can then be processed for RNA-seq (Woronik et al., 2022). A differential expression (DE) analysis of the transcriptomes obtained from pooled tail tips of animals with and without TTM yielded 564 genes that are directly or indirectly controlled by DMD-3 and are candidates for being involved in TTM.

Using gene ontology enrichment analysis and information from the literature, we found groups of DE genes that are acting in TTM and are potentially co-regulated. We call these gene-groups modules. Specifically, we find that chondroitin proteoglycan synthesis, glycerol synthesis and the UPR (unfolded protein response) pathway are activated during TTM, and cuticle maintenance is repressed. There are also 40 transcription factors among the DE genes and severa lintracellular modulators, i.e. GTPases and kinases. We use this information to propose an updated model for the type of regulatory network that might act downstream of DMD-3 in TTM.

## Results and Discussion

### RNA seq of laser-dissected tail tips

To determine which genes are important for morphogenesis of the male tail tip, we sequenced the transcriptome of laser microdissected tail tips of animals at the beginning of the L4 stage when DMD-3 is expressed but TTM had not started yet. Worms were synchronized by L1 arrest. We sampled pools of 230-400 tail tips from 4 different conditions: wild-type (WT) males and *dmd-3(gf)* hermaphrodites that undergo TTM (TTM+), and WT hermaphrodites and *dmd-3(-)* males that do not undergo complete TTM (TTM-). The CEL-Seq2 method (Hashimshony et al., 2016) was used to prepare libraries from extracted RNA of each pool. With this method, we obtained on average ∼653,000 UMI (unique molecular identifiers) per sample (Table S1).

### Principal Component Analysis

To conduct sample level quality control and to explore the strong patterns driving variation in our dataset, we performed a principal component analysis. Visualization of the principal component analysis revealed that all samples except those from *dmd-3(gf)* hermaphrodites clustered by tail tip phenotype (Fig. 2A). *dmd-3(-)* males and WT hermaphrodites do not contain DMD-3 and are TTM-cluster together, while samples of WT male tails, which undergo DMD-3 directed TTM, cluster separately. WT male samples do not cluster as tightly as the *dmd-3(-)* male and WT hermaphrodite samples. This pattern likely arises because the dynamic gene expression exhibited during TTM exacerbated slight variations in developmental timing across samples.

**Figure 2.**
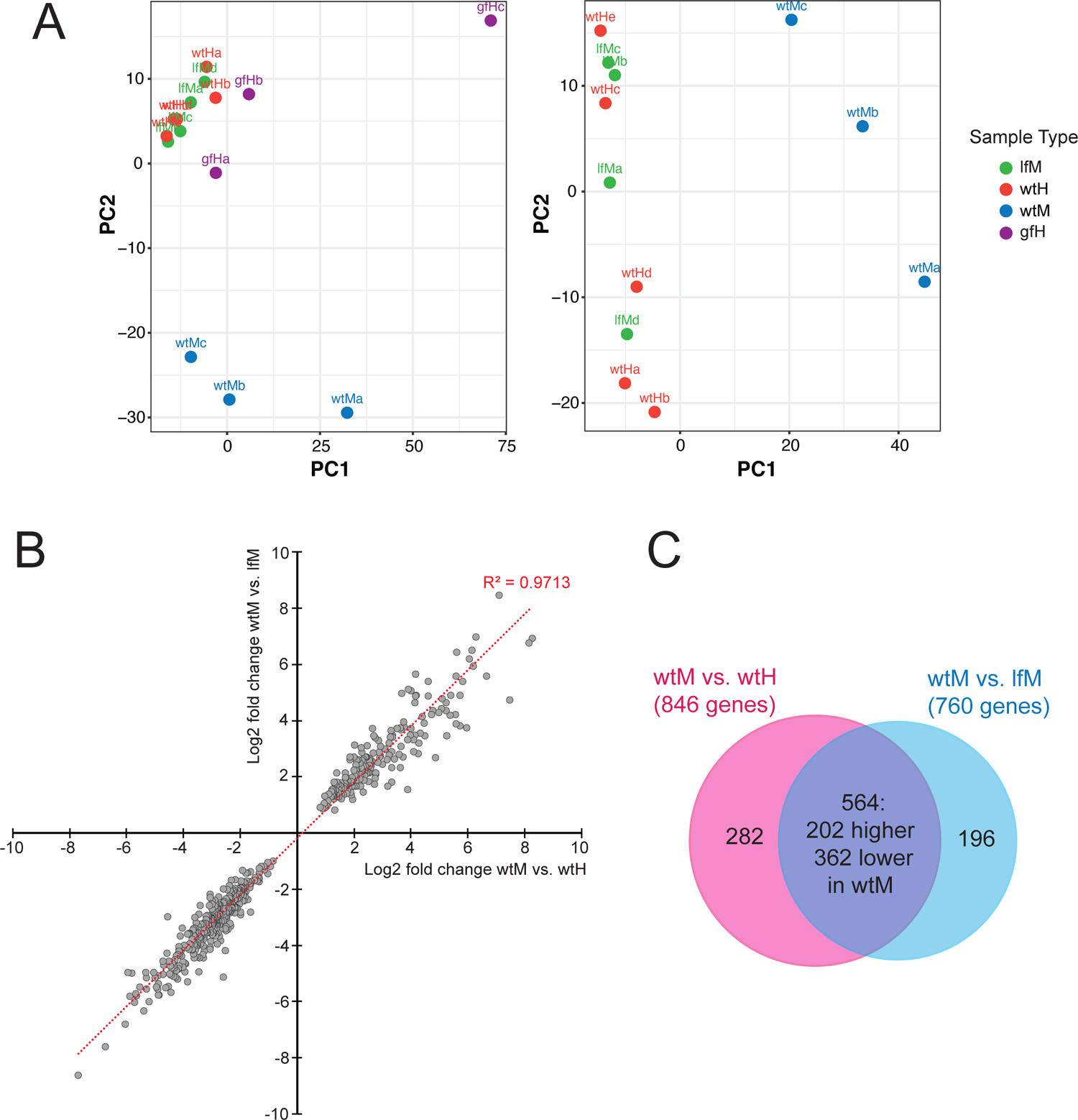
(A) Principal component analysis using all samples, (left) and only samples for WT males (wtM) WT hermaphrodites (wtH) and *dmd-3(-)* males (lfM) excluding *dmd-3(gf)* hermaphrodites (gfH). (B) log_2_ fold changes for each gene in the comparisons of WT males vs. WT hermaphrodites plotted against the log_2_ fold change in the comparison of WT males with *dmd-3(-)* males. (C) Venn diagram showing the overlap of DE genes found in the comparison of WT males with hermaphrodites and WT males with *dmd-3(-)* males.

The *dmd-3(gf)* hermaphrodite bioreplicates do not cluster together in the PCA plot, nor do they cluster with WT male samples as might be expected for tail tips that undergo TTM. These large differences between *dmd-3(gf)* hermaphrodite samples might be due to heterochronic shifts in the development of the tail tip in these mutants: some animals initiate TTM precociously in L3 (Mason et al., 2008). For this reason we did not have confidence in our ability to draw biological conclusions from these samples and excluded them from downstream analyses.

### DE analysis

Next, we conducted a differential expression analysis on two comparisons in our dataset: WT males vs. *dmd-3(-)* males and WT males vs. WT hermaphrodites. Using a cutoff for the adjusted P-value of 0.01, we found 760 genes that were differentially expressed between WT and *dmd-3(-)* males and 846 genes that were differentially expressed between WT males and hermaphrodites. A comparison of the log_2_-fold change of 564 candidate genes that overlap in the two DE analyses showed that not only was the direction in which expression differed the same (e.g. genes have higher expression in in WT males than in hermaphrodites *and* in *dmd-3(-)* males), but also the degree to which these genes were DE (Fig. 2B). The overlap of these gene sets is visualized in Fig. 2C. The 282 genes DE in male vs. hermaphrodite comparison but not in the WT male vs. *dmd-3(-)* male comparison (blue sector) are sex-specific but not regulated by DMD-3, and thus not likely to be important for TTM. The 196 genes that are DE in wtM vs. lfM but not DE in wtM vs. wtH (pink sector) are interpreted to be DMD-3 targets but might not be important for TTM. Thus, we focused primarily on the 564 genes in the intersection (purple); we call these our high quality (HQ) candidates. Of the 564 HQ candidate genes, 202 showed higher expression in WT males (Table S1). Because DMD-3 is only expressed in WT male tail tips, we can conclude that these 202 genes are directly or indirectly activated by DMD-3, and the remaining 362 genes are repressed by DMD-3.

### Validation

#### Comparison with results of an RNAi screen for TTM defects

To validate the candidate TTM genes obtained with our RNA-seq data, we compared the list of 564 high-quality candidate genes with the 210 genes that yielded a TTM phenotype in a previous genome-wide, post-embryonic RNAi feeding screen (Nelson et al., 2011). We found an overlap of only 11 genes (Table 1). RNAi of these genes led to a Lep phenotype (a protruding tail tip in adult males resulting from defective TTM). Seven are activated and four are repressed by DMD-3. Five DE genes were not evaluated by Nelson et al. (2011) because RNAi led to larval lethality. One of these genes is *lin-41*, which was shown to have an over-retracted (Ore) RNAi phenotype (Del Rio-Albrechtsen et al., 2006).

**Table 1:**
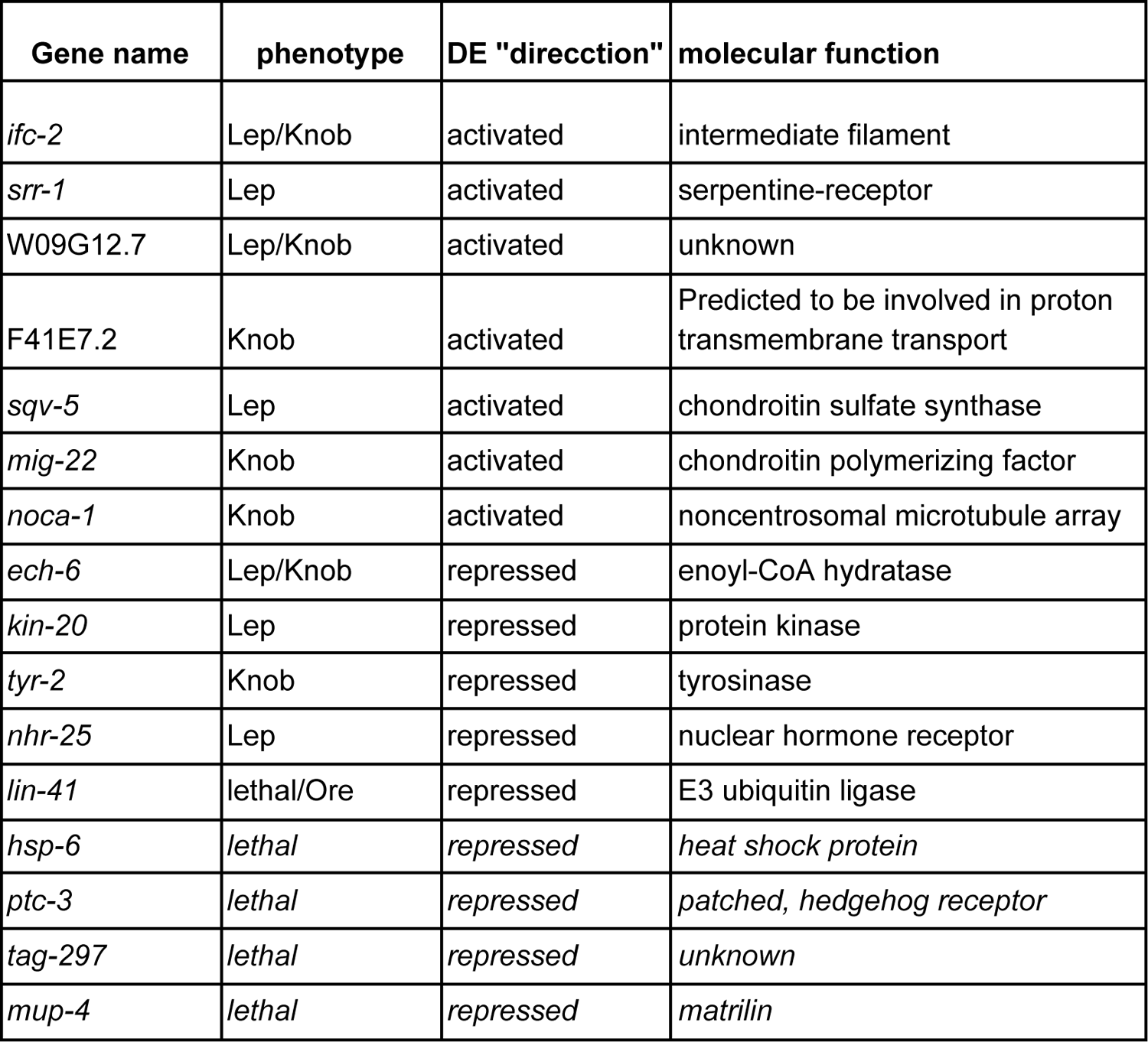
Overlap of the high quality DE genes with genes that showed a phenotype in a whole-genome RNAi screen for TTM defects (Nelson et al., 2011) (Lep = partially unretracted tail tip in adult males, indicating partial failure of TTM, Knob = similar to Lep but smaller).

| Gene name | phenotype | DE "direction" | molecular function |
| --- | --- | --- | --- |
| *ifc-2* | Lep/Knob | activated | intermediate filament |
| *srr-1* | Lep | activated | serpentine-receptor |
| W09G12.7 | Lep/Knob | activated | unknown |
| F41E7.2 | Knob | activated | Predicted to be involved in proton transmembrane transport |
| *sqv-5* | Lep | activated | chondroitin sulfate synthase |
| *mig-22* | Knob | activated | chondroitin polymerizing factor |
| *noca-1* | Knob | activated | noncentrosomal microtubule array |
| *ech-6* | Lep/Knob | repressed | enoyl-CoA hydratase |
| *kin-20* | Lep | repressed | protein kinase |
| *tyr-2* | Knob | repressed | tyrosinase |
| *nhr-25* | Lep | repressed | nuclear hormone receptor |
| *lin-41* | lethal/Ore | repressed | E3 ubiquitin ligase |
| *hsp-6* | *lethal* | *repressed* | *heat shock protein* |
| *ptc-3* | *lethal* | *repressed* | *patched, hedgehog receptor* |
| *tag-297* | *lethal* | *repressed* | *unknown* |
| *mup-4* | *lethal* | *repressed* | *matrilin* |

The small overlap between these two gene lists is not surprising for several reasons. First, the RNAi screen identified genes that were both upstream and downstream of DMD-3 in the TTM gene regulatory network, while our differential expression analysis is expected to only identify genes that are downstream of DMD-3. Second, we know of several genes that are important for the process of TTM, but are likely regulated post-transcriptionally, e.g. *cdc-42* and genes for components of the cytoskeleton where activation or localization rather than changes in expression might be required for TTM. Such genes may show an RNAi phenotype without being DE. Third, we expect a large amount of redundancy in pathways involved in morphogenesis (Sawyer et al., 2011; Wieschaus, 1997). For that reason, many genes that are DE may not show any phenotype when knocked down by RNAi. Indeed, the observed RNAi phenotype is subtle for most genes. For example, we know that RNAi of the fusogen *eff-1*, which is activated by DMD-3, does not show any TTM phenotype (Mason et al., 2008). Fourth, our RNA-seq dataset only represents a single time point during TTM, while the RNAi screen identified genes important for TTM throughout the L4 to adult transition. In addition, a gene required for TTM may be expressed in a tissue adjacent to the tail tip and regulate TTM cell-nonautonomously, as has been shown for *egl-18* (Nelson et al., 2011). Finally, RNAi is more likely to identify genes whose function is required for TTM than those that need to be turned off. Thus, many of the genes that are repressed by DMD-3 might not show an RNAi phenotype. In this context, it is surprising that any of the repressed genes showed a Lep phenotype. Here, one might expect to observe the opposite of failed TTM, i.e. the over-retracted (Ore) phenotype. Genes that nevertheless show a Lep RNAi phenotype may be in a negative feedback loop with DMD-3, as has been demonstrated for *nhr-25*: NHR-25 is required for the onset of DMD-3 expression, but its expression is in turn repressed by DMD-3 (Nelson et al., 2011).

#### Transcriptional reporter assays

As a second method of validation, we made GFP reporters for the proximal promoter regions of a selection of DE genes. Reporters were built to include the region upstream of the gene between the transcription start site and the next gene. We also included the first exon and the first intron, because the first intron of many genes has been shown to contain important regulatory sites (Fuxman Bass et al., 2014). We focused on genes that were activated by DMD-3 and showed a large log_2_-fold change between WT males and females or *dmd-3(-)* males. We used three criteria to evaluate the expression of the reporters: (1) Is the reporter expressed in the male tail tip during TTM? (2) Is the reporter more brightly expressed in the male tail tip—or in more cells—than in the hermaphrodite? (3) When crossed into the *dmd-3(-)* background, is the reporter not expressed or less brightly expressed than in WT males? Of 49 reporters, 26 were expressed in male tail tips more strongly than in tail tips of hermaphrodites (Table 2, Fig. 3). Seventeen of these reporters were investigated in the *dmd-3(-)* genetic background; 16 showed reduced tail tip expression in males lacking functional DMD-3. Ten additional reporters were expressed in tail tips but were equally bright in males and hermaphrodites (Table 2, Fig. 3). It is expected that more subtle differences in the regulation of the genes between sexes or conditions are not always captured with multi-copy extrachromosomal arrays. Thus, reporters for which we did not see a difference between sexes would not invalidate the DE results. In summary, for many of the most differentially expressed genes, transcriptional reporter expression mirrored the RNA-seq results, providing further confidence that our RNA-seq data set reflects *bona fide* DMD-3-regulated genes that are involved in TTM.

**Figure 3.**
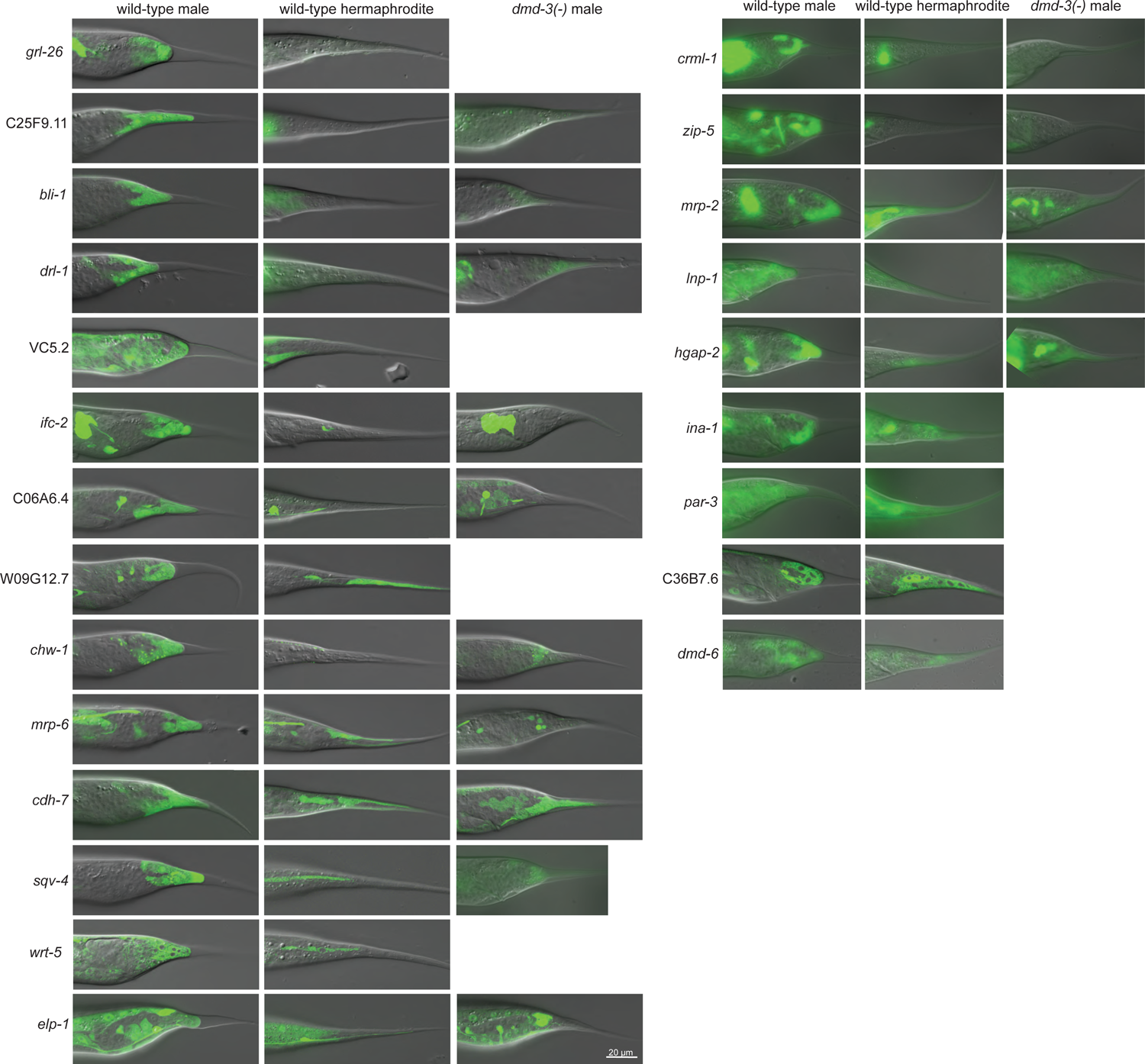
Tail tip expression of transcriptional reporters for DE genes activated by DMD-3 and for *dmd-6*, which is repressed by DMD-3.

**Table 2:** Results of the transcriptional reporter assay. (* = genes repressed by DMD-3)

| gene | expressed in male tail tip | tail tip expression different in hermaphrodites | tail tip expression different in *dmd-3*(-) males | images |
| --- | --- | --- | --- | --- |
| *grl-26* | yes | yes/absent | yes | Fig. 3 |
| C25F9.11 | yes | yes/absent | yes | Fig. 3 |
| *zip-5* | yes | yes/absent | yes | Fig. 3 |
| *bli-1* | yes | yes/absent | yes | Fig. 3 |
| *drl-1* | yes | yes/absent | yes | Fig. 3 |

| **gene** | **expressed in male tail tip** | **tail tip expression different in hermaphrodites** | **tail tip expression different in *dmd-3(-)* males** | **images** |
| --- | --- | --- | --- | --- |
| *ifc-2* | yes | yes/absent | yes | Fig. 3 |
| C06A6.4 | yes | yes/absent | yes | Fig. 3 |
| *chw-1* | yes | yes/absent | yes | Fig. 3 |
| *sqv-4* | yes | yes/absent | yes | Figs. 3, 4 |
| *crml-1* | yes | yes/absent | yes | Fig. 3 |
| *fos-1* | yes | yes/absent | yes | - |
| VC5.2 | yes | yes/absent | no | Fig. 3 |
| *wrt-5* | yes | yes/absent | n.d. | - |
| *phy-2* | yes | yes/absent | n.d. | - |
| *mrp-6* | yes | yes/fewer cells | yes | Fig. 3 |
| *elp-1* | yes | yes/fewer cells | yes | Fig. 3 |
| *mrp-2* | yes | yes/fewer cells | yes | Fig. 3 |
| *cdh-7* | yes | yes/fewer cells | no | Fig. 3 |
| W09G12.7 | yes | yes/fewer cells | n.d. | Fig. 3 |
| *hgap-2* | yes | yes/fainter | yes | Fig. 3 |
| *lnp-1* | yes | yes/fainter | yes | Fig. 3 |
| *mig-22* | yes | yes/fainter | n.d. | - |
| C06E7.88 | yes | yes/fainter | n.d. | - |
| *sqv-5* | yes | yes/fainter | n.d. | Fig. 4 |
| *dpy-6* | yes | yes/fainter | n.d. | - |
| *best-1* | yes | yes/fainter | n.d. | - |
| *srw-85* | yes | no | n.d. | - |
| *sqv-7* | yes | no | n.d. |  |
| *ina-1* | yes | no | n.d. | Fig. 3 |
| *par-3* | yes | no | n.d. | Fig. 3 |
| C36B7.6 | yes | no | n.d. | Fig. 3 |
| *dpm-3* | yes | no | n.d. | - |
| F14E5.2 | yes | no | n.d. | - |
| *dbn-1* | yes | no | n.d. | - |
| F48F7.3 | RnP cells and neurons | - | n.d. | - |
| *vang-1* | in neurons | - | n.d. | - |

| gene | expressed in male tail tip | tail tip expression different in hermaphrodites | tail tip expression different in *dmd-3(-)* males | images |
| --- | --- | --- | --- | --- |
| *ptr-9* | in neurons | - | n.d. | - |
| *mctp-1* | in neurons | - | n.d. | - |
| *rap-1* | in phasmid neuron | - | n.d. | - |
| *col-20* | no | - | n.d. | - |
| R01E6.5 | no | - | n.d. | - |
| Y39B6A.9 | no | - | n.d. | - |
| *aqp-8* | no | - | n.d. | - |
| *lec-6* | no | - | n.d. | - |
| *nex-2* | no | - | n.d. | - |
| tag-294 | no | - | n.d. | - |
| *dmd-6\** | yes | no | n.d. | Fig. 3 |
| F07F6.4* | yes | no | n.d. | - |
| *ccpp-1\** | no | - | n.d. | - |

#### Loss of Function phenotypes

To begin determining which of the DMD-3 regulated genes are essential for TTM, we examined the loss-of-function male tail phenotype for a subset of genes whose transcriptional reporters show clear male-biased expression in the tail tip. To do so, we examined either RNAi treated males (*sqv-4, sqv-5,* and *par-3*) or males homozygous for mutant alleles (*fos-1, grl-26, chw-1, C06A6.4, crml-1, zip-5, bli-1, mrp-2, cdh-7,wrt-5, elp-1,* and *srw-85*). Both *sqv-4* and *sqv-5* RNAi yielded the male tail phenotype that is typical of loss-of-function conditions for genes involved in chondroitin biosynthesis (see below). In contrast, loss-of-function of all other genes examined resulted in adult males with wild type tails. This suggests that the gene network regulated by DMD-3 is robust, with a large degree of functional redundancy and/or multiple pathways with small effects. This is consistent with the results of the RNAi feeding screen which found typically relatively mild defects in TTM (Nelson, et al., 2009). The examination of loss-of-function conditions for additional DMD-3 regulated genes is needed to verify this model.

#### Enrichment analysis

To find biological functions preferentially expressed in the tail tip during TTM, we searched for Gene Ontology (GO) terms statistically overrepresented (at a q value threshold = 0.03) among all DE genes with an adjusted P-value < 0.01. We found that more informative results were obtained when we analyzed the genes activated by DMD-3 separately from the genes repressed by DMD-3 (Table S3).

#### Over-represented GO terms for genes activated by DMD-3

For genes activated by DMD-3, the most highly enriched GO terms were “IRE1-mediated unfolded protein response” (GO:0036498) and “response to topologically incorrect protein” (GO:0035966). These GO terms refer to the UPR (unfolded protein response) which senses and responds to an overabundance of unfolded proteins in the ER. Such proteins are found during normal development in tissues with high protein secretory load (Mitra and Ryoo, 2019). This suggests that the male tail tip during TTM could be an active secretory tissue. However, the UPR also has other noncanonical roles on differentiation and morphogenesis (Hetz et al., 2020). Other notable GO categories in this class of genes are ABC-type transporter activity (GO:0140359) and spindle localization (GO:0051653). The latter GO term is here associated with genes that are involved in cell polarity establishment and cell migration (*hmr-1, par-3, ced-1, ced-6*), processes known to be important for directing morphogenetic events like TTM (e.g. Nance and Priess, 2002; Vuong-Brender et al., 2016).

#### Over-represented GO terms for genes repressed by DMD-3

For the genes repressed by DMD-3, the most highly enriched GO categories are “structural constituent of cuticle” (GO:0042302) with 25 genes for collagens and the cuticlin CUT-2, “collagen trimer” (GO:0005581) and “molting cycle” (GO:0042303). Among the DE genes in the category “molting cycle” are several genes for astacin-like proteases, which are also in the GO category “metalloendopeptidase activity” (GO:0004222) that is enriched as well. Taken together, these GO categories indicate that cuticular collagen synthesis and processing is downregulated in tail tips undergoing TTM.

During TTM, the tail tip cells are largely internalized such that they have only a very small cuticle-covered apical surface in adults. We thus initially assumed that collagens whose genes are repressed by DMD-3 are part of the adult cuticle. To test this idea, we compared our data with those by Abete-Luzi et al. (2020), who identified genes regulated by LIN-29, a transcription factor expressed at the juvenile-to-adult transition and required for several developmental events during this time, including cuticle synthesis and molting. The authors found 33 collagen genes upregulated by LIN-29 and concluded that these collagens are used in the synthesis of the adult cuticle. Only 4 of the LIN-29 upregulated collagen genes are DE in the tail tip, two (*col-77* and *col-99*) are repressed and two (*bli-1* and *col-20*) are activated by DMD-3. Thus, most of the collagens repressed by DMD-3 are likely not part of the adult cuticle. Also, in our experiments, half of the LIN-29 regulated collagens were not or only lowly expressed in the hermaphrodite tail tip, suggesting that our sample time was too early to catch mRNAs of most adult collagens. We thus conclude that the collagens and their modifying enzymes that are expressed in hermaphrodite and *dmd-3(-)* male tails but less so in wild-type male tails are required for maintaining the L4 cuticle. Since the male tail tip cells dissociate from the cuticle at the beginning of L4, it is conceivable that they do not participate in L4 cuticle maintenance. Alternatively, some of these genes could be involved in the synthesis of the provisional apical ECM that precedes the formation of the adult cuticle (Katz et al., 2022).

### Proteins synthesized in the tail tip during TTM

From the GO analysis we concluded that the male tail tip during TTM has a higher protein secretory load than tail tips that do not undergo TTM. Do our data indicate what kind of proteins are produced in the ER during TTM?

#### Tail tip-expressed collagens

Several genes in the GO category “IRE1-mediated unfolded protein response” encode protein disulfide isomerases (PDI): *pdi-1*, *pdi-2*, *pdi-6* and *dnj-27*. Two further paralogs of these genes, *pdi-3* and *C14B9.2* are DE in the comparison of WT and *dmd-3(-)* males, the former narrowly missing our cutoff for the adjusted P-value of 0.01. PDI-1, PDI-2 and PDI-3 were shown to act synergistically in collagen biogenesis (Winter et al., 2007). PDI-2 forms the beta subunits of collagen prolyl 4-hydroxylase (P4H) which is involved in an essential early step of collagen biosynthesis. PDI-2 combines with two α subunits that can be DPY-18 or PHY-2. The gene for PHY-2 is also among our DE genes activated by DMD-3. It is thus possible that these disulfide isomerases are involved in synthesis of specific collagens in the male tail tip during TTM. Our list of 564 DE genes contains 29 collagen genes. However, as noted above, all but 4 are repressed in tail tips undergoing TTM. If collagen biosynthesis is upregulated in the WT male tail tip, it might be for the production of the remaining 4 collagens, BLI-1, COL-20, COL-46 and COL-89.

We decided to study the expression of these upregulated collagens. We therefore tagged the endogenous BLI-1, COL-20 and COL-89 N-terminally with fluorescent proteins, and investigated an integrated multicopy reporter for BLI-1 C-terminally tagged with GFP. Unexpectedly, BLI-1 and COL-20 are localized in the cytoplasm of the tail tip cells and not external to the cells (Fig. 4). Expression of BLI-1 in hyp8-11 and hyp13 starts right after the L3/L4 molt and continues into adulthood. Beginning at stage L4.4 (for stage assignment see Kiontke et al. submitted), BLI-1::GFP is also expressed in the body epidermis in hermaphrodites and males, where it is later localized to the struts of the adult cuticle as previously described (Johnson et al., 2023; Lints and Hall, 2005). The N-terminally tagged BLI-1 reporter is expressed in the hypodermis in mid-L4 after which hypodermal expression disappears, probably because GFP is cleaved off at the RXXR furin cleavage site along with the N-terminal portion of the protein (Johnson et al., 2023). Tail tip cell expression, however, is still seen in young adult males. The COL-20 protein begins to be expressed in the seam of both sexes late during the L4/adult lethargus phase. In early adults, the GFP::COL-20 reporter is seen in hyp7 (Fig. 4). This reporter is not expressed in the hermaphrodite tail tip cells or in the vulva. In males, GFP::COL-20 is first expressed at stage L4.2 in only one cell, likely P11.ppap, which is part of the hook sensillum. In stage L4.8 and L4.9 males, expression is also seen in another cell in the hook, possibly P11.ppp and in a cell in the posterior part of the proctodeum, which might be B.pap or B.paa. At this stage, seam expression also begins. In young adult males, GFP::COL-20 is expressed in the cytoplasm of the ventral tail hyp as it migrates anteriorly. It is also seen in cells of the proctodeum, hyp7 and possibly hyp11. GFP::COL-89 shows very bright expression in cells of the male proctodeum which largely conceals the much fainter tail tip expression when observed with a wide-field microscope. Only one cell in the tail (likely R9.p) occasionally expresses COL-89 more brightly (Fig. 4). In hermaphrodites, GFP::COL-89 is seen in cells of the proctodeum and in the vulval cells of the vulB2, vulC and vulD toroids.

**Figure 4.**
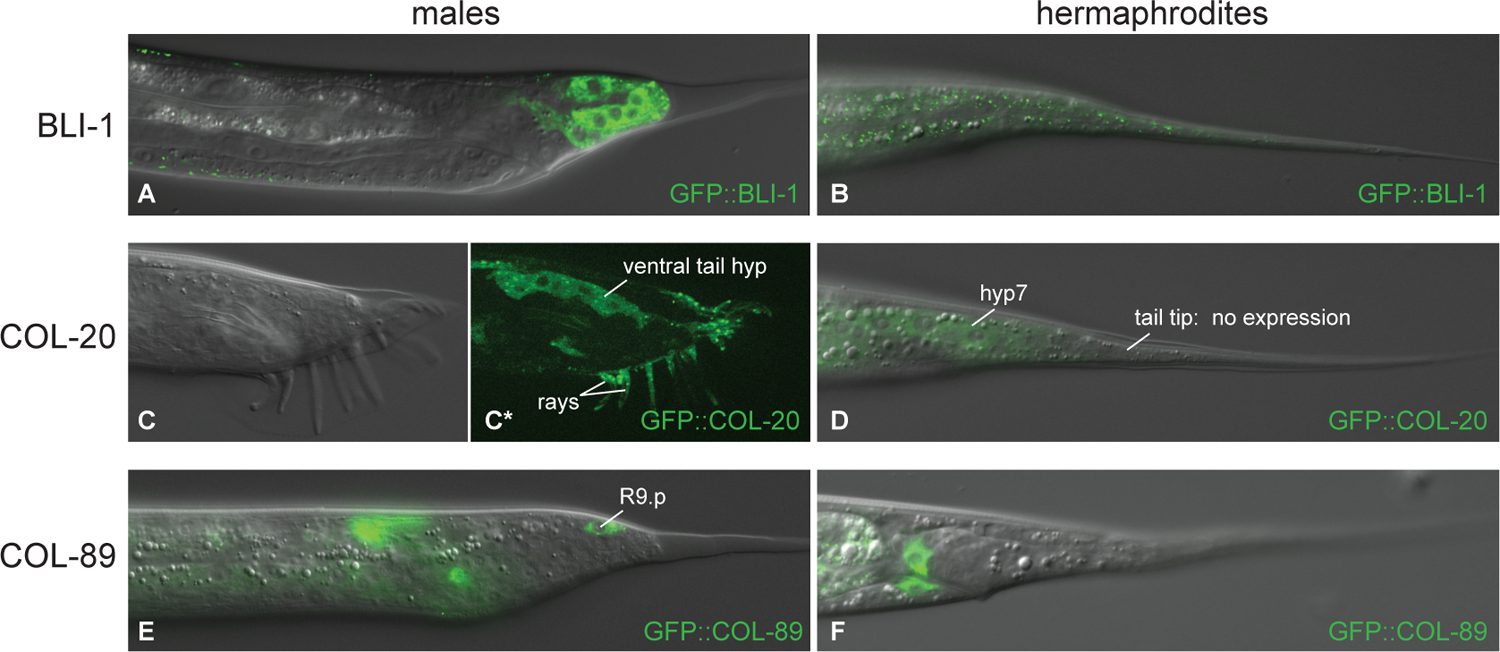
Expression of three collagens that are activated by DMD-3. (A, B) GFP::BLI-1 expression in an L4.4 male (A) and L4.5 hermaphrodite (B). In males, BLI-1 is expressed in the tail tip cells and in hyp13 and weakly in puncta at the apical surface of hypodermal cells, where it is also observed in hermaphrodites. (C, D) GFP::COL-20 begins in late L4 in the seam and hyp7 in both sexes but not in the hermaphrodite tail tip. In young adult males (C, C*), COL-20 is expressed in the ventral tail hyp and the rays. (E, F) GFP::COL-89 is expressed in the proctodeum of both sexes and occasionally in R9.p in males.

It appears that all three collagens are expressed in the tail tip cells of males but not hermaphrodites. As a caveat, we do not know where the cleaved COL-20 and COL-89 are located since our reporters were N-terminally tagged and the fluorescent protein is cleaved off during collagen processing.

#### Chondroitin proteoglycan synthesis

Several genes in the chondroitin synthesis pathway are upregulated in the tail tips of males undergoing TTM. Genes in this pathway were initially identified in a screen for mutations in vulva morphogenesis that resulted in the “Sqashed vulva” (Sqv) phenotype (Herman et al., 1999). Subsequently, the *sqv*-genes were shown to encode transporters and enzymes that are required for the production of chondroitin proteoglycans, proteins to which a polysaccharide chain is attached (reviewed in Berninsone, 2006). The chondroitin synthesis pathway comprises 8 SQV proteins and the chondroitin polymerizing factor MIG-22 (Suzuki et al., 2006). Genes for SQV-4, SQV-5 SQV-7 and MIG-22 are DE in tail tips with higher expression in wt males. An additional nucleotide sugar transporter gene, *nstp-4*, is also DE, as well as the uncharacterized gene C29H12.2 with a putative sugar phosphate transporter domain. We analyzed transcriptional reporters for *sqv-4*, *sqv-5* and *sqv-7* and found that all three are expressed in tail tips (Fig. 5). Except for *sqv-4*, the reporters were expressed in tails of males and hermaphrodites. RNAi-knockdowns of the genes for SQV-5 and for MIG-22 are known to cause Lep tail tip phenotypes (Nelson et al. 2011), highlighting the importance of this pathway for TTM.

**Figure 5.**
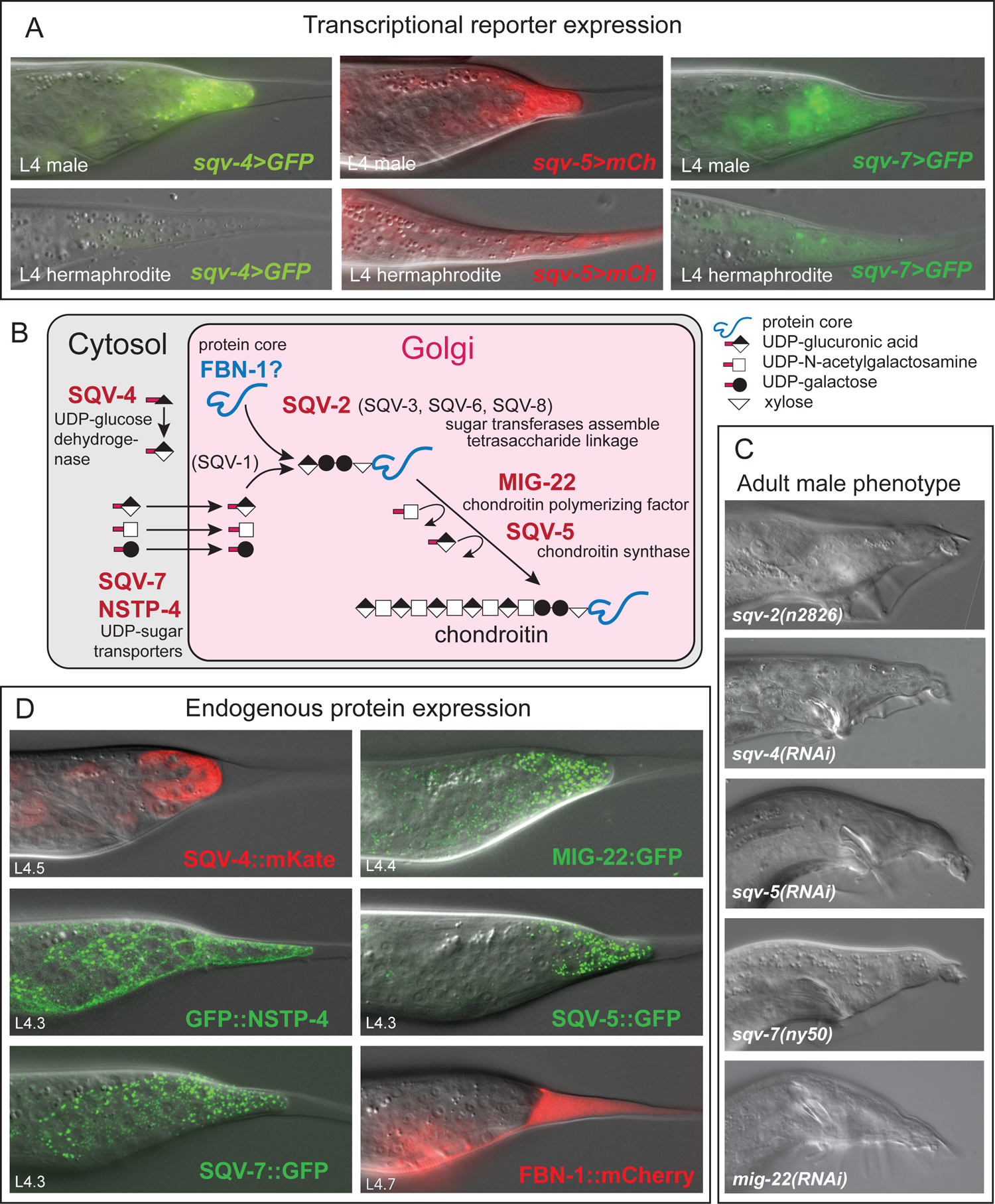
The chondroitin synthesis pathway plays a role in TTM. (A) Expression of transcriptional reporters *sqv-4, sqv-5* and *sqv-7* during L4 in males and hermaphrodites. (B) Schematic of the pathway (modified after Hwang et al. 2003) with genes found to be involved in TTM in red. (C) Examples of the mutant or RNAi tail tip phenotype for 4 *sqv-*genes and *mig-22*. (D) Examples for the protein expression pattern in the L4 male tail tip of 5 components of the *sqv-*pathway and the putative core protein FBN-1.

To confirm these results and to investigate if knockdown or mutation of other *sqv*-genes would also show a tail tip phenotype, we performed RNAi by feeding or injection and observed male tails of mutants. We found that mutation or RNAi of *sqv-2*, *sqv-4* and *sqv-5* led to similar defects in the morphology of the adult male tail (Fig. 5C) with a slightly unretracted tail tip (and defects in further anterior tail structures). In the course of tagging the endogenous *sqv-7* gene with GFP using CRISPR/Cas9 genome editing, we observed several F1 males with Lep tails, indicating that here, the function of this gene was impaired. This mutation could not be isolated, probably because it causes sterility or embryonic lethality.

To further investigate the expression of the proteins in the *sqv*-pathway during TTM, we tagged endogenous SQV-4, SQV-5, SQV-7, MIG-22 and NSTP-4 with GFPnovo2 or mKate, respectively. SQV-4::mKate is expressed cytoplasmically as expected (Berninsone, 2006). All other proteins are seen in puncta, consistent with their localization to the Golgi apparatus (Fig. 5D). In the tail tip, SQV-4::mKate expression begins after the L3/L4 molt in all tail tip cells and hyp13 and shows a marked increase as TTM progresses (Fig. S1). In contrast, SQV-4::mKate expression is faint in hermaphrodite tails tips and does not increase during L4. When crossed into the *dmd-3(-)* background, brightening of the reporter is delayed (Fig. S1). SQV-5::GFP expression is seen in male tail tips in bright puncta (Fig. 5D). In hermaphrodite tail tips, SQV-5::GFP is also present, but expression is less bright and the puncta are smaller (Fig. S2). Expression of the reporter is present in the tail tip of *dmd-3(-)* males, but it is less bright than in wild-type males (Fig. S2). SQV-7::GFP is expressed in the tail tip and the Rn.p cells as well as the seam throughout L4 with increasing brightness in tail tip (Fig. 5D), the seam and tail seam as development progresses. In hermaphrodites, SQV-7::GFP is expressed in the seam and weakly in the tail tip (data not shown). A GFP::NSTP-4 reporter is expressed in the tail tip and in other cells in the developing tails, where it is concentrated at the membranes between the cells in puncta and diffuse in the cytoplasm (Fig. 5D). It is also expressed less brightly in hyp7 throughout L4. In hermaphrodites, GFP::NSTP-4 is seen in the vulva cells, the gonad, hyp7 and the seam. MIG-22::GFP is seen in hyp7 and the tail tip in males and hermaphrodites. However, in males, tail tip expression is brighter than expression in other tissues. The reporter is also seen in hyp13 (not shown).

Although the chondroitin synthesis pathway was elucidated by studying vulva morphogenesis mutants, the chondroitin core proteins in the developing vulva remain unknown. It is likely that more than one core protein is involved because no single-gene mutation causes a vulva defect (Hwang and Horvitz, 2002). Using mass spectrometry, Noborn et al. (2018) identified many of the chondroitin core proteins (CPGs); currently 23 are known and two more are predicted. We detected mRNA for 11 of these proteins in tail tips. Only *fbn-1* and the predicted CPG B0365.9 are DE in the comparison of WT males vs. WT hermaphrodites, but neither is among our HQ DE genes. A translational reporter for *fbn-1* is expressed in the tail of L4 males, first only in the proctodeum and the area around the anus in the extracellular space between tail tissue and L4 cuticle. Beginning with stage L4.4, FBN-1::mCherry is also seen in the extracellular space behind the tail tip where expression gets brighter as TTM continues. In L4.9 males, FBN-1 is only seen in the region of the developing fan but not inside of the L4 tail tip cuticle. It is thus conceivable that in the male tail, FBN-1 is one chondroitin proteoglycan core protein that is secreted as the tail tip cells retract. FBN-1 also plays a role in vulva development. Here, this protein is laid down around a central core, which is proposed to also contain chondroitin proteoglycans (Cohen et al., 2020). FBN-1 appears in the vulva beginning with stage L4.2 with peak expression during stages L4.5 and L4.6, the same time when its expression is brightest in the male tail. Given that FBN-1::mCherry expression appeared relatively late during TTM, it is likely that other chondroitin proteoglycans are being secreted at an earlier stage, similar to those forming the central core in the developing vulva (Cohen et al., 2020).

#### Transcription factors downstream of DMD-3

Previous work identified a number of transcription factors (TF) involved in TTM (Nelson et al., 2011). Most of these were hypothesized to act upstream of DMD-3, like the HOX genes *nob-1* and *php-3* and GATA factor genes *egl-18* and *elt-6*. We sought to identify additional TFs that act downstream of DMD-3 by searching our list of high-confidence DE genes for TFs listed by Ma et al. (2021) and the database cisBP (Weirauch et al., 2014), and other putative TFs described in WormBase. We found 40 genes likely involved in transcriptional control (Table 3), 11 of them activated and 28 repressed by DMD-3. Among the latter is *nhr-25*, which was shown to be repressed by DMD-3 (Nelson et al., 2011). Another known TTM TF is TLP-1 (Zhao et al., 2002). Its gene is repressed by DMD-3, even though its mutant phenotype is Lep. It is possible that TLP-1, like NHR-25 acts in a negative feedback loop with DMD-3, first activating its expression and then being repressed by DMD-3 (Nelson et al., 2011).

**Table 3:** Transcription factors and post-translational modifiers among 564 DE genes.

| activated by DMD-3 |  | repressed by DMD-3 |  |
| --- | --- | --- | --- |
| gene name | Gene-stable ID | gene name | Gene-stable ID |
| TFs |  | TFs |  |
| *egl-46* | WBGene00001210 | *die-1* | WBGene00000995 |
| *fos-1* | WBGene00001345 | *egl-13* | WBGene00001182 |
| *zip-5* | WBGene00007932 | *fkh-9* | WBGene00001441 |
| *jun-1* | WBGene00012005 | *mxl-3* | WBGene00003511 |
| *dmd-3* | WBGene00012832 | *nhr-17* | WBGene00003616 |
| *bed-2* | WBGene00012943 | *nhr-25* | WBGene00003623 |
| *ztf-22* | WBGene00012988 | *nhr-31* | WBGene00003625 |
| *tag-294* | WBGene00016937 | *nhr-45* | WBGene00003635 |
| M03D4.4 | WBGene00019751 | *nhr-46* | WBGene00003636 |
| *tsct-1* | WBGene00011824 | *nhr-88* | WBGene00003678 |
| C06A6.2 | WBGene00015507 | *nhr-97* | WBGene00003687 |
| *jmjd-3.1* | WBGene00017571 | *pat-9* | WBGene00003933 |
| Kinases |  | *sem-2* | WBGene00004771 |
| *drl-1* | WBGene00017578 | *sem-4* | WBGene00004773 |
| *gck-1* | WBGene00001526 | *sex-1* | WBGene00004786 |
| *gck-2* | WBGene00022603 | *sox-3* | WBGene00004950 |
| *wnk-1* | WBGene00006941 | *tlp-1* | WBGene00006580 |
| Phosphatases |  | *dmd-6* | WBGene00007058 |
| *ptp-1* | WBGene00004213 | C01G6.5 | WBGene00007227 |
| *ptp-3* | WBGene00004215 | *bed-3* | WBGene00009133 |
| GTPases |  | *ccch-1* | WBGene00009532 |
| *chw-1* | WBGene00009059 | *atfs-1* | WBGene00013878 |
| *ral-1* | WBGene00021811 | *bcl-11* | WBGene00017430 |
| *rap-1* | WBGene00004307 | *hmbx-1* | WBGene00018786 |
| GAPs |  | *dmd-7* | WBGene00019521 |
| *tbc-17* | WBGene00012894 | *ztf-16* | WBGene00019960 |
| *hgap-2* | WBGene00008430 | *ell-1* | WBGene00021281 |
|  |  | *dve-1* | WBGene00022861 |
|  |  | Kinases |  |
|  |  | *kin-20* | WBGene00002203 |
|  |  | *nsy-1* | WBGene00003822 |
|  |  | *pps-1* | WBGene00004091 |
|  |  | *prk-2* | WBGene00004183 |
|  |  | R06A10.4 | WBGene00019911 |
|  |  | *sek-1* | WBGene00004758 |
|  |  | *sma-5* | WBGene00004859 |
|  |  | *tpa-1* | WBGene00006599 |
|  |  | Phosphatases |  |
|  |  | *ipp-5* | WBGene00002146 |
|  |  | *pgph-3* | WBGene00016892 |
|  |  | *scpl-3* | WBGene00021629 |
|  |  | T13H5.1 | WBGene00011755 |
|  |  | Y62E10A.13 | WBGene00013379 |
|  |  | phosphatase regulator |  |
|  |  | T01D1.3 | WBGene00020148 |
|  |  | GTPases |  |
|  |  | *rap-3* | WBGene00004309 |
|  |  | *gpa-12* | WBGene00001674 |
|  |  | GAP |  |
|  |  | F07F6.4 | WBGene00017217 |
|  |  | GEFs |  |
|  |  | *rei-1* | WBGene00007270 |
|  |  | F55D12.2 | WBGene00010111 |

To further investigate possible connections between the TFs downstream of DMD-3, we turned to information about TF targets in the database TFlink (Liska et al., 2022) and the TF yeast-2-hybrid data by Reece-Hoyes et al. (2013) (Fig. S2). This analysis showed that many but not all of the TFs downstream of DMD-3 are targets of the well-studied TFs JUN-1 and FOS-1 (both activated by DMD-3) and/or NHR-25 (repressed by DMD-3), which is a DNA-binding scaffold protein that requires binding of other factors for specific gene regulatory capacity (Asahina et al., 2020). The most protein interactions among the TFs downstream of DMD-3 are with EGL-13, a TF that in is involved in fate specifications of neurons and uterine cells (Feng et al., 2013; Gramstrup Petersen et al., 2013; Oommen and Newman, 2007).

### Post-translational modifiers downstream of DMD-3

Cellular functions are regulated not only by transcription factors but also by kinases, phosphatases and GTPases. We found several genes for such modifiers to be DE in our analysis (Table 3). Among the kinases repressed by DMD-3 is *kin-20*, one of the few DE genes that also shows an RNAi phenotype. Two kinases, DRL-1 and WNK-1, whose expression is activated by DMD-3, were found to act in the *C. elegans* hypertonic stress response (HTSR). Both activate the transcription of *gpdh-1*, which is a rate-limiting enzyme for glycerol synthesis (Urso and Lamitina, 2021; Wimberly and Choe, 2022). *drl-1* is specifically expressed in tail tip cells during TTM. Among the DE GPTases is *chw-1* with a male-specific expression pattern. CHW-1 is known from its involvement in polarity establishment during vulva development, where it boosts signaling through the WNT receptor LIN-17/Frizzled (Kidd et al., 2015).

### Hedgehog-like signaling

Another group of notable DE genes in TTM are those in Hedgehog-like signaling. The *C. elegans* Hedgehog (Hh) signaling pathway is divergent from the well-studied pathway in other organisms with absence of clear orthologs of the Hh ligand and the signal transduction molecule SMO (Zugasti et al., 2005). However, there are many Hh related molecules as well as an expansion of receptors related to patched (Zugasti et al., 2005). Fourteen of 64 Hh-related putative ligands are DE, all but three repressed by DMD-3. The activated Hh-related genes are *grl-26* (with one of the largest log_2_-fold changes in our dataset) *grd-16*, and *wrt-5*. Of the 27 patched and patched-related genes, 10 are DE. Only one of them, *ptr-9*, is activated by DMD-3. Hh signaling is essential for development. In *C. elegans*, it has been implicated e.g. in molting (Lažetić and Fay, 2017), vulva development (Zugasti et al., 2005) and in the organization of the precuticular ECM (Cohen et al., 2021). Zugasti et al. (2005) also found male tail morphogenesis defects upon RNAi of 6 ptr-genes. Three of them, *ptr-18*, *ptr-20,* and *ptr-23* are repressed by DMD-3 in the tail tip, suggesting that they may function in the tissue anterior of the tail tip.

### DMD-3-repressed genes with unknown function

Among the genes repressed by DMD-3 with the smallest log2-fold change in our DE analyses are many with unknown function. Some of these genes are highly expressed in TTM-tail tips but either not expressed at all in TTM+ tails, or less by an order of magnitude. For a subset of these genes, interactions were found or predicted previously (interaction information in WormBase). We used this information to generate a network and found interaction clusters for 117 of the DMD-3-repressed genes (Fig. S3, Table S6). Most of the genes with unknown functions cluster together with the cuticlin gene *cut-2* (one of the most highly expressed genes in the TTM-tail tips), the zinc carboxypeptidase *suro-1* known to be involved in cuticle synthesis (Kim et al., 2011) and two patched-related receptors, *ptr-4* and *ptr-16*, also known to be involved in cuticle formation (Cohen et al., 2021) (cluster 1 in Fig. S3). This cluster is linked to two other clusters that contain primarily collagens. We thus hypothesize that the 36 uncharacterized genes in cluster 1 function in the context of cuticle maintenance or synthesis.

## Conclusion

Through tail-tip-specific RNA-seq, we identified 564 genes that are differentially expressed in WT male tail tips that express DMD-3 and undergo TTM vs. tail tips that do not express DMD-3 and do not undergo TTM (hermaphrodite and *dmd-3(-)* male tails). We verified several candidates by male-tail specific transcriptional reporter expression. Analysis of the DE genes suggests that the male tail tip has high protein production during TTM but downregulates cuticle maintenance. Forty transcription factors and 30 post-translational modifiers are DE.

### Regulatory interactions downstream of the core transcription factor DMD-3

After analyzing the transcriptome of the tail tip undergoing TTM and finding genes that are under the direct or indirect transcriptional control of DMD-3, are we closer to understanding the architecture of the gene network downstream of DMD-3 in this morphogenetic process?

Morphogenesis involves universal and pleiotropically acting “cellular machines” that are specialized for particular aspects of cell behavior (e.g. apicobasal polarity, actomyosin, endocytosis, cell-cell and cell-ECM adhesion). How the activity of these machines is integrated in a particular morphogenetic event is dependent on cues from the cellular environment, which are relayed by tissue specific transcription factors. However, transcription factors are unlikely to affect these pleiotropic cellular machines directly, but instead regulate components of signaling pathways, such as kinases and G-proteins or their GEF or GAP switches (Bernadskaya and Christiaen, 2016), which we here call “effectors”. In addition, we posit that morphogenesis employs other groups of proteins that act together in various contexts but are not universally present in every cell. We call such groups of proteins “conditional modules”. These conditional modules may also be regulated by effectors rather than by transcription factors. Under these assumptions, one model for the TTM system architecture could be quite simple, with few direct DMD-3 targets, all of which are effectors (Fig. 6A). This network model is similar to that for *Ciona* cardiac progenitor migration where the transcription factors Mesp and FoxF control cellular processes through receptors for signaling pathways, Rho GTPase and aPKC (Bernadskaya and Christiaen, 2016). An alternative model is that DMD-3 targets parallel cascades of transcription factors, which—likely also through effectors—regulate TTM (Fig. 6B). Vertebrate neural crest formation is an example for such a regulatory architecture (Bernadskaya and Christiaen, 2016). We are only at the beginning of evaluating which type of network model is more likely to be true for TTM. However, the observation that 40 transcription factors are DE in the tail tip during TTM, strongly suggests that transcriptional control plays a larger role in TTM than, for instance, in *Ciona* cardiac progenitor migration. At the same time, the TTM transcriptome also contains genes for a number of GTPases, their regulators, kinases, and phosphatases to be DE. Therefore, our current working hypothesis for the TTM network model is one where DMD-3 regulates different modules through a variety of transcription factors in addition to regulating effectors that facilitate the appropriate activity of universal pleiotropic cellular machines Fig. 6C).

**Figure 6.**
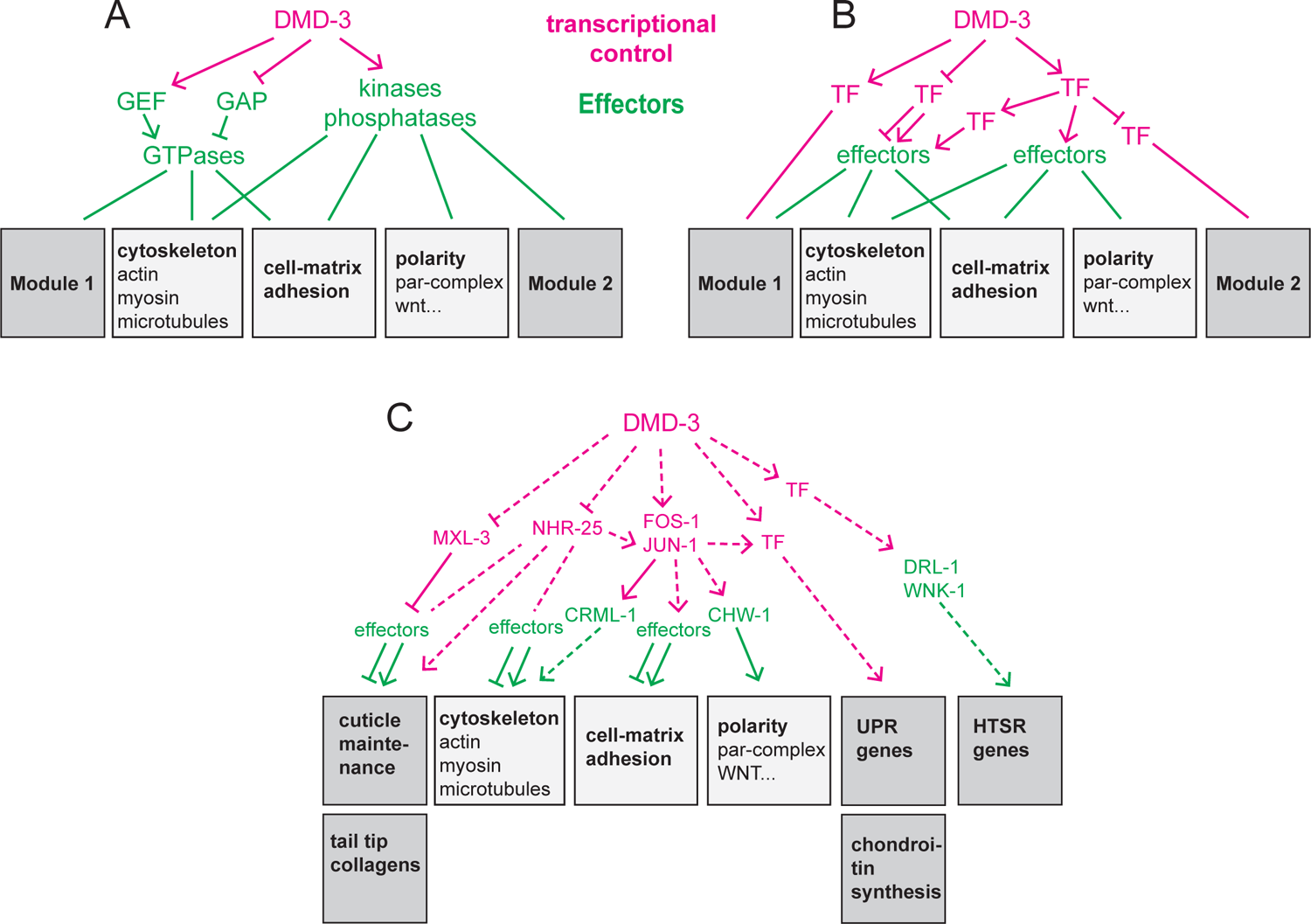
Network models for TTM. (A) DMD-3 controls only few targets, all of them GEFs, GAPs, kinases and phosphatases (effectors). (B) Between DMD-3 and the effectors or conditional modules are chains of transcription factors. (C) Composite model with a combination of transcription factors and effectors controlling molecular machines and conditional modules.

### Conditional modules in TTM

Analysis of GO terms and previous knowledge of the genes DE in the tail tip during TTM revealed several groups of genes that likely act together in sub-processes of TTM, and are thus candidates for members of specific conditional modules.

#### Genes that respond to incorrectly folded proteins

UPR (unfolded protein response) pathway components are enriched among the genes activated by DMD-3. The UPR is also activated in the anchor cell prior to invasion. Here, ribosome biogenesis is also upregulated to accommodate increased translation of transmembrane and secreted proteins (Costa et al., 2023). Although we do not find enrichment of ribosome biogenesis genes in TTM, we believe that the UPR is activated due to increased translation in the tail tip as well.

#### Genes for maintaining the L4 cuticle

GO enrichment analysis found among the genes repressed by DMD-3 an overabundance of genes for cuticle proteins, cuticle development or the molting process. While these genes are employed during each molting cycle with peak expression just prior to or during the molt (Hendriks et al., 2014), they are repressed in the male tail tip during TTM. We assume that these cuticle proteins and their regulators form a module required for cuticle maintenance in tail tips that do not undergo TTM. We hypothesize that the genes of unknown function in interaction cluster 1 (Fig. S3) also belong to this module. This module is turned off in WT males.

#### Genes for the male tail tip expressed collagens

BLI-1, COL-20, COL-89, and COL-46 and the collagen processing proteins PHY-2 (proline hydroxylase), PDI-1, PDI-2 (Protein Disulfide Isomerases), FKB-5 and FKB-7 (FK506-Binding proteins) (Page and Johnstone, 2007) are all activated by DMD-3. *pdi-1* and *pdi-2* are targets of the transcription factor MXL-3 according to TFlink (Liska et al., 2022), which is repressed by DMD-3. We found BLI-1 and COL-20 in the cytoplasm of the tail tip cells. The role of this module is unclear. However, it should be noted that three of the collagens (*bli-1, col-20* and *col-89*) were found in an RNAi screen for genes that affect the expression of *gpdh-1.* Signaling through collagens (*dpy-7, −8, −9,* and *-10)* affects *gpdh-1* expression (represses it). It is thus possible that BLI-1, COL-20, and COL-89 also signal to the glycerol synthesis pathway, promoting *gpdh-1* expression.

#### Genes involved in cell fusion

Most cell fusions in *C. elegans* are mediated by the fusogen EFF-1 (Podbilewicz, 2006), the gene for which is DE/activated by DMD-3. Because gap junctions were found in the male tail and the developing excretory system along cell borders prior to fusion, it was suggested that communications across these junctions may be important for fusion (Hall, 2017). In the male tail, RNAi against the gap junction genes *inx-12* and *inx-13* cause the Lep phenotype, and *inx-13* is DE/activated in the comparison of wtM vs. lfM. It is possible that *eff-1* and these innexins act together in cell fusion during TTM. Peak expression of transcriptional reporters for *inx-12* and *inx-13* is observed late during TTM when fusion of the tail tip cells is already ongoing (Nelson et al., 2011). Thus, the innexins may have additional functions independent of fusion.

#### Genes in the chondroitin proteoglycan synthesis pathway

(Fig. 5) are activated by DMD-3. The synthesis of chondroitin proteoglycans was shown to be required for vulva development (Berninsone, 2006) and during the assembly of the eggshell (Olson et al., 2012). We show that this module is also deployed during TTM.

#### Genes in the hypertonic stress response (HTSP)

When *C. elegans* encounters high salinity, it elicits the HTSP, which leads to the intracellular accumulation of the osmolyte glycerol. Glycerol synthesis is upregulated by transcriptional activation of the glycerol-3-phosphate-dehydrogenase GPDH-1 (reviewed in Urso and Lamitina, 2021) in part through the kinases, DRL-1 and WNK-1. *gpdh-1, drl-1* and *wnk-1* are DE/activated by DMD-3 in the tail tip, indicating that here, too, glycerol synthesis is upregulated. In HTSP, *gpdh-1* expression is also affected by signaling through the patched-related transmembrane protein PTR-23 (Rohlfing et al., 2011); RNAi against its mRNA leads to male tail morphogenesis and molting defects (Zugasti et al., 2005). In the tail tip, DMD-3 represses *ptr-23*. Previous RNAi screens for genes affecting the expression of *gpdh-1* transcriptional reporters (Chandler et al., 2023; Rohlfing et al., 2011) found several genes that are also DE in the tail tip: *atx-2, bbln-1, bli-1, col-20, col-89, crt-1,* F01G10.9, *ndg-4, srw-85* (all activated by DMD-3), *col-77, ell-1* F01G10.10, *fath-1, gdh-1,* H03E18.1, *hum-5, moe-3, pgph-3 and ptr-4* (repressed by DMD-3). Also, 18 of the genes with a *gpdh-1* RNAi phenotype were also positive in the RNAi screen for TTM defects (Nelson et al., 2011). Among these is the gene for the tail-tip-expressed WNT LIN-44 and the gene for the GEF VAV-1, which was previously shown to be regulated by DMD-3 (Nelson et al., 2011). Finally, two collagens, *dpy-7* and *dpy-9*, which are both involved in HTSR signaling (Dodd et al., 2018), also showed a TTM RNAi phenotype. This overlap of RNAi-positives for HTSR and TTM suggests an overlap of the function of these genes as well. Another DE gene activated by DMD-3 and involved in the HTSP is that for the aquaporin AQP-8. Aquaporins are channel-forming proteins that allow for the passage of water and small molecules through cell membranes. *aqp-8* mRNA levels increase approximately eightfold during hypertonic stress (Huang et al., 2007). Finally, *lea-1*, a gene that is upregulated during desiccation stress (Hibshman and Goldstein, 2021) is DE in the comparison of wtM vs. lfM. Taking all of this together, we hypothesize that the HTSP genes, *aqp-8* and possibly *lea-1* constitute a module in TTM.

We propose that the module for chondroitin proteoglycan synthesis and the HTSR module function together in TTM (Fig. 7): For vulva development, it was proposed that the hygroscopic chondroitin sulfate draws water into the vulva lumen, thereby expanding it and creating pressure that prevents the vulva from collapsing (Hwang et al., 2003). It is now clear that chondroitin proteoglycans act with other aECM proteins in a complex luminal scaffold to shape the vulva lumen during morphogenesis. However, the initial lumen appears to be created by chondroitin proteoglycans (Cohen et al., 2020). We propose that likewise, in TTM, the extracellular space is filled with a hygroscopic matrix as the tail tip tissue dissociates from the L4 cuticle and rounds up. We further hypothesize that chondroitin proteoglycans draw water from the tail tip tissue into the extracellular space rather than from the environment, leading to hypertonic stress in the tail tip cells. To counteract this stress, *gpdh-1* transcription and osmolyte production are activated.

**Figure 7.**
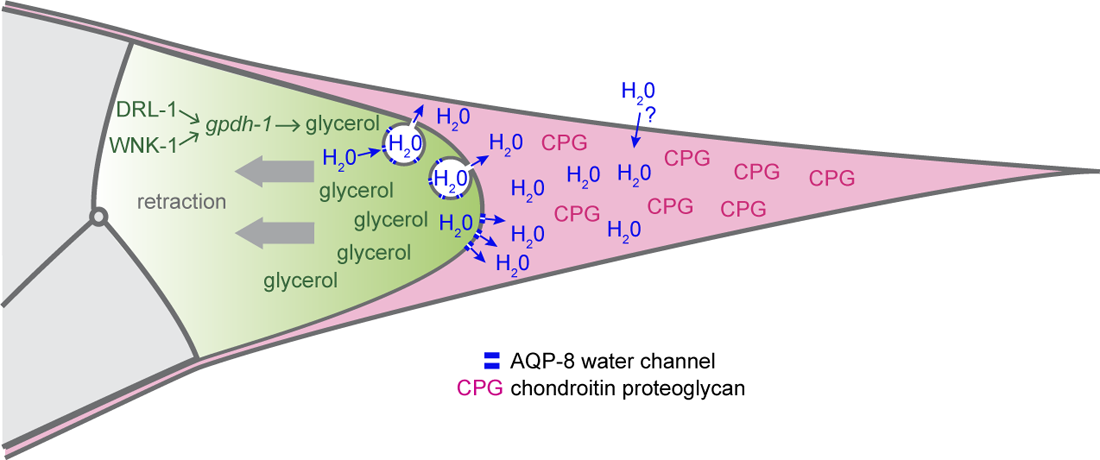
Hypothetical role of chondroitin proteoglycans and components of the HTSR in TTTM. The tail tip secretes CPGs into the extracellular space between tail tip tissue and cuticle (pink). This hygroscopic matrix attracts water, which passes through aquaporin-8 water channels that are located in the membrane of the tail tip or in vesicles similar to those constituting the canaliculi of the excretory duct. The thus created pressure aids/stabilizes tail tip cell shortening and retraction. The osmolarity in the tail tip tissue is balanced by glycerol production through upregulation of gpdh-1 activity via the kinases DRL-1 and WNK-1.

Consistent with this model, the water channel protein AQP-8 is also activated by DMD-3. In HTSR, *aqp-8* transcription and translation are elevated upon exposure to high osmolarity in the environment, probably to allow increased excretion through the excretory canal (Igual Gil et al., 2017). The flux of water into the lumen of the excretory canal is regulated by the number of water channels that are located in canaliculi, chains of vesicles continuous with the canal lumen. If water transport from tail tip cells into the extracellular space were similar to that in the excretory canal, we would expect to find vesicles in the cytoplasm of tail tip cells undergoing TTM. Such vesicles were observed in one male by Nguyen et al. (1999), although the authors interpreted them as the remnants of membranes between fused cells rather than vesicles transporting water and other cargo to the tail tip. Testing this model and evaluating how genes in other modules function in TTM and how they are regulated by DMD-3 will be a subject of future research.

## Materials and methods

### Worm husbandry and handling

*C. elegans* strains were grown on a lawn of *E. coli* (OP50-1) according to standard methods (Lewis and Fleming, 1995; Stiernagle, 2006).

### Synchronization

To obtain synchronized populations of larvae, two methods were used: L1 arrest and “hatch-off”. For L1 arrest (Lewis and Fleming, 1995), gravid hermaphrodites were treated with alkaline hypochlorite to release embryos. Embryos were then washed with M9 buffer and left shaking in M9 buffer at 25°C for 18-24 hours. Arrested L1’s were placed on plates seeded with *E.coli* and allowed to develop at 25 °C until the desired developmental time. For hatch-off, the method described in Woronik et al. (2022) was used. Briefly, gravid hermaphrodites were allowed to lay eggs overnight. Then, hermaphrodites and all hatched larvae were removed by washes with M9 buffer. The remaining embryos were incubated at 25 °C. After one hour, all L1 that had hatched during this time were collected in M9, placed on food and allowed to develop to the L3 stage (∼20-22 h post hatch at 25 °C), at which time males could be distinguished from hermaphrodite and picked onto separate plates.

### Strains used

Table S7.

### Tail-tip-specific RNA-seq

This experiment was performed with four types of samples: tail tips of L4 wild-type hermaphrodites (N2), wild type males (CB4088), *dmd-3*(-) males (UR156) and *dmd-3(gf)* hermaphrodites (UR257).

### Collecting synchronized L4 males or hermaphrodites

Worm development was synchronized by L1 arrest: Gravid CB4088 hermaphrodites were exposed to an alkaline hypochlorite solution (Porta-de-la-Riva et al., 2012) to release embryos. Embryos were then washed with M9 buffer and left shaking in M9 buffer at 25°C for 18-24 hours. Arrested L1s were placed on plates seeded with *E.coli* and allowed to develop at 25 °C for 22 hours after which time L3 males could be distinguished from hermaphrodites. Males were picked onto a new plate under a dissection microscope at room temperature. To collect hermaphrodites, the N2 strain was used, but L4 animals were still picked to account for imperfect synchronization. After 2 hours of picking, worms were collected into a tube with M9 buffer and washed twice with cold M9 buffer supplemented with a small amount of detergent (Tween20). After washing, the worms were fixed in ice cold freshly made 70% methanol, washed twice in 70% methanol and stored in this solution for at least 1 hour and at most 24 hours until further processing.

### Laser microdissection and RNA extraction

Drops of worms fixed in 70% methanol were pipetted onto a Leica PEN membrane slide (76 x 26 mm P/N 11505158). The slide was then placed on a slide warmer at 50°C to evaporate the methanol. Once dried, the slide was mounted onto a Leica laser dissection microscope. Costar Thermowell tubes (0.5 ml with flat cap, Cat#: 6530) were loaded into the collection holder, and 65µL of lysis buffer (Ambion RNAqueous-Micro Kit, ThermoFisher Scientific, cat # AM1931) was pipetted into the tube cap. Using the 40x objective, 50-100 tail tips were dissected into each cap. Before unloading the tube from the collection holder, an additional 35µL of lysis buffer was added to the tube cap. Tubes were spun down, flash frozen in liquid nitrogen or on dry ice and stored at −80 °C until enough tissue was collected for one RNA extraction (240-440 tail tips).

RNA was extracted using the Ambion RNAqueous-Micro kit following the manufacturer’s protocol and eluted twice into 8 µl elution solution. The quality and quantity of the extracted RNA was evaluated with Agilent high sensitivity RNA screen tapes (#5067-5579).

### Library preparation and sequencing

To obtain enough of DNA for sequencing, the single cell RNA-Seq method CEL-Seq2 (Hashimshony et al., 2016) was used for library construction. Instead of single cells, purified RNA was used as starting material for the protocol, which was otherwise followed as published. For the CEL-Seq2 method, RNA of each sample is indexed at the first reverse transcription step and UMIs are introduced. Subsequently, several samples are combined into the same library.

Here, two libraries were made with the same sample RNAs at two starting concentrations: 300 pg/sample and 118 pg/sample. Each library contained four replicate samples for *dmd-3(-)* males, three for wild-type males, and five for wild-type hermaphrodites (Table 1). The CEL-Seq2 method includes a linear and a PCR amplification step and preserves strandedness: briefly, RNA is reverse transcribed with oligo-dT primers. After second strand synthesis, the DNA is in-vitro transcribed and thereby amplified. The amplified RNA is quantified, fragmented and reverse transcribed with random hexamers. cDNA that contains adapter sequences introduced during the first cDNA synthesis step are subsequently PCR amplified (11 cycles). During this step, Illumina sequencing adapters are added. This method results in 3’ reads that can be easily mapped back to the *C. elegans* genome. Non-polyadenylated RNAs and splice variants are not captured with this method. Libraries were sequenced on two lanes of an Illumina HiSeq 2500 in Rapid Mode (2 x 50, 15 x 36 bp) by the Genomics Core at New York University’s Center for Genomics and Systems Biology.

### Analysis

Raw reads were adapter trimmed using CutAdapt (v1.12) (Kechin et al., 2017). Fastqc (v. 0.11.8) (LaMar, 2015) was used to evaluate read quality. Trimmed reads were mapped to the *C. elegans* genome (WB235) using Bowtie2 (v. 2.3.2) (Langmead and Salzberg, 2012) via the CelSeq2 analysis pipeline (v. 0.2.5; FASTQ_QUAL_MIN_OF_BC: 10, CUT_LENGTH: 35, ALN_QUAL_MIN: 0) (Hashimshony et al., 2016). The read counts matrix generated by the CelSeq2 pipeline was input for an analysis in DESeq2 (v. 1.26.0) (Love et al., 2014). Genes with no mapped reads were removed from the matrix, then differential expression analyses (alpha = 0.01, p adjusted < 0.01) were conducted for wild-type versus *dmd-3*(-) males, wild-type males versus wild-type hermaphrodites, *dmd-3*(-) males versus wild-type hermaphrodites and wild-type hermaphrodites versus *dmd-3(gf)* hermaphrodites. We then identified genes that were differentially expressed in both TTM(+) versus TTM(-) comparisons (i.e. wild-type versus *dmd-3*(-) males, wild-type males versus wild-type hermaphrodites and *dmd-3(gf)* hermaphrodites versus wild-type hermaphrodites), but not in the TTM(-) versus TTM(-) comparison (i.e. *dmd-3*(-) males versus wild-type hermaphrodites) to generate a list of high-quality candidate genes involved in TTM.

To gain biological insights for our high-quality candidates, we performed enrichment analyses using the Enrichment Analysis Tool (q value threshold = 0.1) on Wormbase (v. WS290) (Angeles-Albores et al., 2016; Angeles-Albores et al., 2018). The Enrichment Analysis Tool utilizes a hypergeometric statistical model to identify gene, anatomy, and worm phenotype ontology terms that are over-represented in a list of genes.

### Transcriptional reporters (Table S8 and S9)

Transcriptional reporters were made either by overlap extension PCR (Nelson and Fitch, 2011) or by Gateway cloning, restriction enzyme cloning or Gibson assembly (Gibson et al., 2009).

Generally, a region extending from the gene upstream of the transcription start site of a candidate gene plus the first exon and intron of that gene was fused to the sequence of GFP in plasmid pPD95.75 (Fire Vector kit, Addgene) which also contains the *unc-54* 3’UTR. For PCR, Q5 high fidelity DNA polymerase (NEB), PrimeStar Max (Takara) or PrimeStar GXL (Takara) were used. The PCR products were purified with the Promega Wizard SV Gel and PCR clean-up system or the Qiaquick PCR purification kit (Qiagen). For restriction enzyme cloning, the regulatory region of each gene was amplified and ligated into *SphI/AgeI* or S*phI/KpnI* digested pPD95.75. For Gateway cloning, the regulatory region of *sqv-5* was amplified, inserted into a pENTRY vector, and transferred into pENTRY-*SrfI*-mCherry using the Multisite Gateway System (ThermoScientific). For Gibson Assembly, the regulatory region of each gene was inserted into *SmaI* digested pUC19 together with GFP amplified from pPD95.75 using NEBuilder® HiFi DNA Assembly Master Mix (NEB - E2621). PCR products or Plasmids were micro-injected into CB4088 [him-5(e1490)] hermaphrodites with myo-2::mCherry from plasmid pCFJ90 (Addgene) as co-injection marker, or into EM574 [*pha-1(e2123); him-5(e1490)*] with plasmid pBX as co-injection marker, which rescues the temperature sensitive lethal phenotype of the *pha-1(e2123)* mutation. Microinjections were performed as previously described (Evans, 2006), several lines were maintained and examined for GFP expression in males and hermaphrodites with emphasis on expression in the tail tip. Some reporters were then crossed into strains containing the *dmd-3(-)* allele.

### Translational reporters (Table S8)

For some candidates we made endogenously tagged translational reporters via CRISPR Cas9 genome editing. To identify PAM sites near the 3’ or 5’ end of a gene, we used the algorithms implemented in Benchling (https://www.benchling.com). If a suitable PAM site was within 0-3 nucleotides of the insertion site (before translation start or stop), we made homology repair templates with 35 nucleotide long homology arms. For PAM sites further away from the insertion site, homology repair templates were designed with 200 nucleotide long homology arms. Oligonucleotides with these overhangs and a region complementary to the sequence of GFPnovo2 (in plasmid pSM Addgene) or mKate (in plasmid pCFJ350 Addgene) were used to PCR amplify the homology repair template (Takara PrimeStar Max). After purification (Promega Wizard SV Gel and PCR clean-up system), the concentration of the HRT was adjusted to 100 ng/µl as measured with Nanodrop (Thermo Scientific). Guide RNAs were purchased from IDT either as sgRNA or as crRNA and resuspended in IDT Duplex buffer to a concentration of 100 µM or 200 µM, respectively. crRNAs were annealed to the IDT tracerRNA (200 µM) at 95 °C for 5 minutes. The concentration of the guide RNA was adjusted with water to 62 µM. 0.5 µl of IDT Cas9 protein was mixed with 0.5 µl of guide RNA and incubated for 5 minutes at room temperature. The HRT was heated to 95 °C for 5 minutes and immediately placed on ice. An injection mix was made by adding to 1 µl RNP complex 1 µl 1xTE buffer (pH8), 2 µl HRT and 6 µl water. The mixture was loaded into needles and injected into both gonads of 10 young hermaphrodites. No coCRISPR reagents were used. 16-32 L4 stage progeny of the injected hermaphrodites were picked onto individual plates and kept at 25 °C until they had laid several eggs. Then, each F1 hermaphrodite was picked into 10 µl proteinase K lysis buffer (Fay and Bender, 2006), frozen in liquid nitrogen, thawed and refrozen 3 times and incubated for 90 minutes at 60 °C and 15 minutes at 95 °C. This worm lysis was used for PCR with primers flanking the insertion site. L4 hermaphrodite progeny of worms with edits were picked onto individual plates and allowed to lay eggs. Lysis and PCR were performed, and this process repeated until a homozygous animal was identified.

### Mutating the *him-5* and *dmd-3* locus with CRISPR Cas9 genome editing

To introduce the *him-5(e1490)* mutation into some strains, we turned to CRISPR Cas9 genome editing, which was performed as described above with a homology repair template that was ordered as ssDNA from IDT. Individual F1 hermaphrodites were isolated as L4 and allowed to reproduce. From plates that contained males, L4 hermaphrodites were picked to individual plates to homozygoze the mutation. No PCR test for an edit was performed.

To create a *dmd-3(null)* mutation, we deleted the entire coding region of the gene with two CRISPR Cas9 cuts and repaired it with a homology repair template that bridged the gap. Homozygous mutants were isolated as for *him-5* above by observing the Lep phenotype in the F1 progeny.

### Microscopy and image processing

For microscopy, animals were either picked from mixed staged plates or from bleach synchronized populations. Worms were paralyzed in 20 mM sodium azide and mounted on a pad of 4 % Noble Agar in 50 % M9 buffer supplemented with 20 mM sodium azide. They were examined with a Zeiss AxioImager equipped with Colibri LED illumination and an Apotome.

Generally, image stacks were recorded with 0.5 µm distance between slices. If the fluorescent signal was too dim for Apotome imaging, the deconvolution algorithm implemented in the Zeiss ZenBlue software v. 2.6 was used (“good/fast iterative” setting) to improve image quality. Some DIC and fluorescent images were generated using a Nikon Ti-S compound inverted microscope equipped with a Hamamatsu C11440-10C Flash 2.8 camera. The NIS Elements Version 4.2 software was used for image acquisition, and digital images were processed using the Adobe Photoshop Elements software (Adobe Systems, San Jose, CA).

### RNAi

RNA interference was performed by injection or feeding. For *sqv-4(RNAi)* by injection, the *sqv-4* coding sequence (+23 to +1397) with flanking T7 promoters was amplified from *C. elegans* RNA using SuperScript One-Step RT-PCR with Platinum Taq (Invitrogen). This PCR product was used as a transcription template to generate *in vitro* synthesized double stranded RNA using the MEGAscript RNAi Kit (Ambion). Purified double-stranded RNA (100 ng/μl) was injected into the gonads of *rrf-3(pk1426) II; qIs56 [lag-2::GFP] him-5(e1490) V* adult hermaphrodites and F_1_ adult males were examined for tail tip phenotypes. *sqv-5* and *par-3* RNAi feeding was carried out as previously described (Kamath et al., 2001) using the pPD129.36 nhr-67 RNAi construct from the Ahringer Library (Geneservice). Briefly, HT115(DE3) bacterial cells were freshly transformed with either the pPD129.36 *sqv-5* or *par-3* RNAi plasmid. Plasmid-containing bacteria were induced to express double-stranded RNA using IPTG and plated on M9 lactose plates. *rrf-3(pk1426) II; qIs56 [lag-2::GFP] him-5(e1490) V* worms were bleach synchronized, and L1-arrested worms were plated onto RNAi-expressing bacteria. The tail phenotype of adult males was assessed ∼3 days later. To generate males in strains with wildtype alleles of *him-5*, worms were grown on bacteria expressing double stranded RNA for *him-8/klp-16* using the pLT 651 plasmid (Addgene plasmid # 59998, Timmons et al., 2014).

### Network analysis and visualization

Interaction data of repressed genes was mined from WormBase using their REST API (http://rest.wormbase.org/rest/field/gene/|WORMBASE_GENE_ID|/interactions). We did not discriminate in the type of interaction, i.e. predicted, physical, regulatory, or genetic. Interactions between DMD-3 repressed genes and non-repressed genes were removed. As the final network is undirected, interactions that differed only in their directionality were collapsed into one. The AutoAnnotate app (version 1.4.1) (Kucera et al., 2016) in Cytoscape was used to calculate clusters using the MCL clustering algorithm (Van Dongen, 2008).

## Acknowledgements

We thank the ∼50 undergraduate Siena College and 6 NYU students who helped generate transcriptional reporters for DMD-3 regulated genes. We specifically thank independent research students Liam Peterson, Chelsea LeBlanc, George Bushey, Athena Bennani, and Lillian Gardner and the students in Adam Mason’s 2020, 2022 and 2023 Developmental Biology Course. We further thank Shehana Gunasekara for help with the CELSeq2 protocol and Raza Mahmood for help with data analysis. Some strains were provided by the CGC, which is funded by NIH Office of Research Infrastructure Programs (P40 OD010440). This work was funded by NSF grant 1656736 and NIH grant R01GM141395 to David H. A. Fitch and NSF grant IOS-1255877 to D. Adam Mason.

## Data availability

The sequencing reads are available at the SRA database under BioProject ID PRJNA1046486. Supplemental tables and figures will be submitted to figshare. Strains are available upon request from the authors; some may be submitted to the Caenorhabditis Genetic Center.

