## Supplemental Figure 1 for "Tissue-specific RNA-seq defines genes governing male tail tip morphogenesis in *C. elegans*"

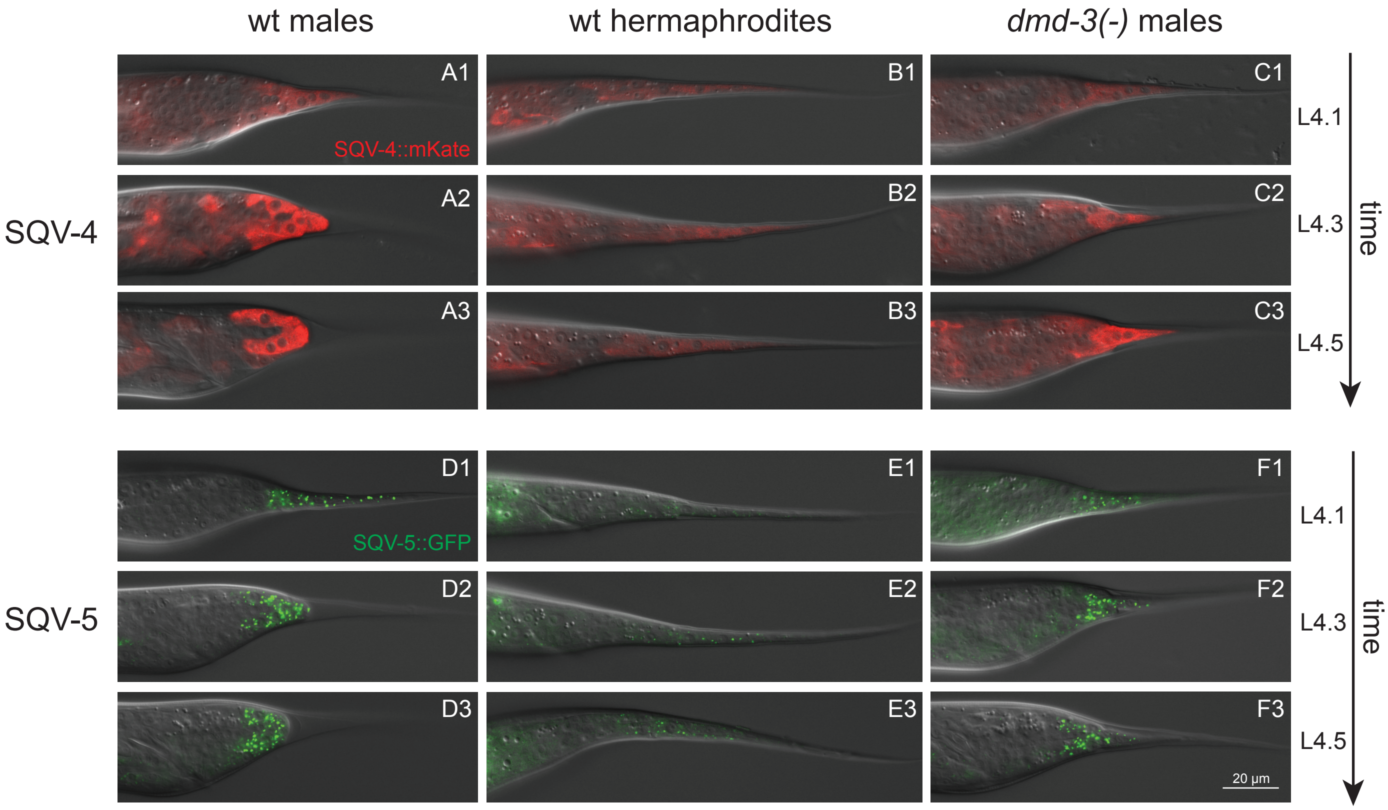


**Figure S1**
Expression of endogenous SQV-4 tagged with mKate (top 9 images) and SQV-5 tagged with GFPnovo2 (bottom 9 images) in tails at three time points (1 = L4.1, 2 = L4.4, 3 = L4.6) during L4 development in WT males (A and D), WT hermaphrodites (B and E) and *dmd-3(-)* males (C and F). Column A: In WT males, SQV-4 is weakly expressed early in L4. Expression increases through stages L4.3-6. Column B: In hermaphrodites, expression is present but faint throughout L4. Column C: in *dmd-3(-)* males, an increase in expression is noticeable but it is delayed relative to WT males. Column D: SQV-5 expression is visible as puncta in WT male tail tip cells throughout L4. Column E: In hermaphrodites, expression is weakly seen in small puncta. Column F: In *dmd-3(-)* male tail tips, SQV-5::GFP expression is seen throughout TTM but it is less intense than in WT males.
