## Supplemental Figure 2 for "Tissue-specific RNA-seq defines genes governing male tail tip morphogenesis in *C. elegans*"

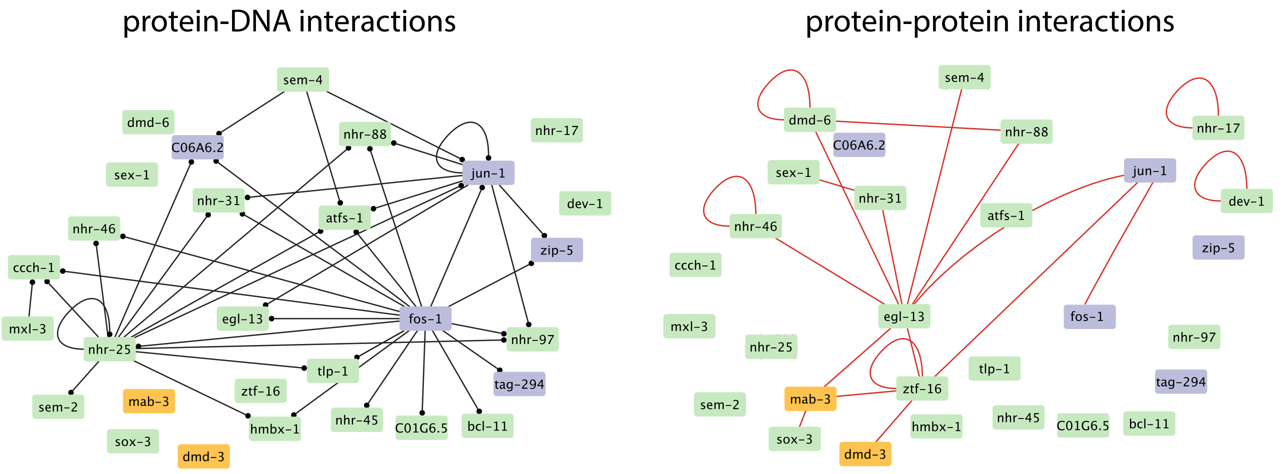


**Figure S2**
TF-target interactions (from TFlink) and protein-protein interactions (Reece-Hoyes et al., 2013) between TFs that are regulated by DMD-3 (DMD-3-activated genes in purple, DMD-3-repressed genes in green).
