## Supplemental Figure 3 for "Tissue-specific RNA-seq defines genes governing male tail tip morphogenesis in *C. elegans*"

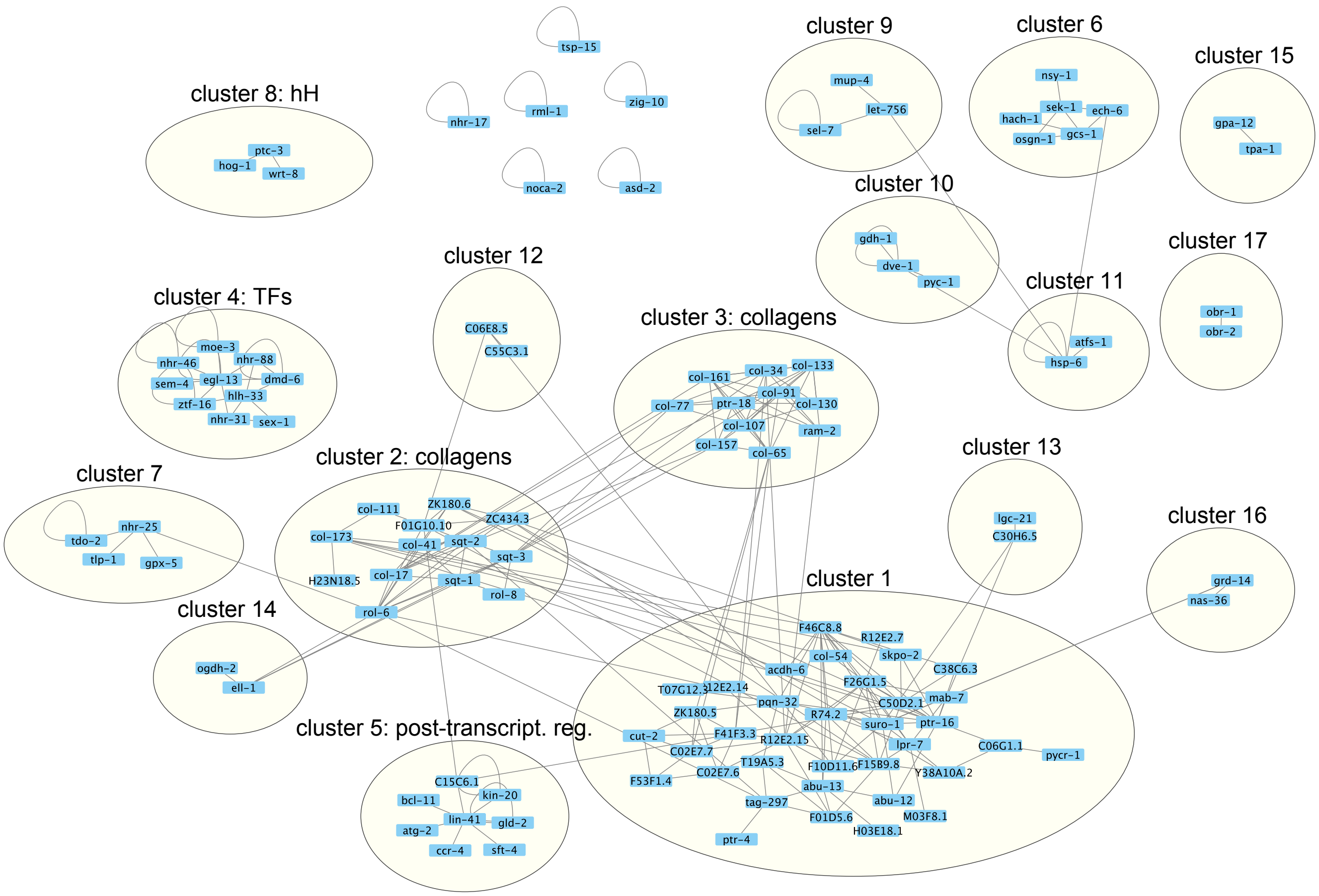


**Figure S3**

Clustering of genes repressed by DMD-3 by interactions mined from WormBase. Interaction cluster 1 contains 20 genes of unknown function.
